# Cardiac fibroblasts counterbalance cardiomyocytes in *LMNA* cardiomyopathy pathogenesis

**DOI:** 10.1101/2025.06.05.657412

**Authors:** Kunal Sikder, Elizabeth Phillips, Nesrine Bouhrira, David Mothy, Nadan Wang, Gisèle Bonne, Kenneth B. Margulies, Jason C. Choi

## Abstract

Genetic cardiomyopathies arising from mutations in the *LMNA* gene, encoding nuclear intermediate filaments lamin A/C, display variable age of onset, severity, and fibrosis development. This variability suggests a fundamental element in disease pathogenesis that has yet to be elucidated. Given the central role cardiac fibroblasts play in fibrosis, we explored the relevance of lamin A/C in cardiac fibroblast function, as very little is known in this regard. Using primary cardiac fibroblasts and *in vivo* mouse models, we show that *Lmna* mutations impact various aspects of cardiac fibroblast function in response to myocyte damage. We show that both lamin A/C depletion and point-mutant variant expression impair cardiac fibroblast proliferation and contraction whereas other functions such as cell migration appears to be mutation dependent. *In vivo* depletion of lamin A/C simultaneously in cardiomyocytes and cardiac fibroblasts significantly delayed disease progression, improved cardiac function, and prolonged survival, indicating that lamin A/C mediate an opposing balance between cardiomyocytes and cardiac fibroblasts in driving disease pathogenesis. Our results elucidate previously unexplored roles of lamin A/C in cardiac fibroblasts and suggest that interactions between cardiac fibroblasts and cardiomyocytes are important determinants of the rate of progression and the severity of *LMNA* cardiomyopathy.

## Introduction

A-type lamins A and C (lamin A/C) are nuclear-resident proteins belonging to type V intermediate filament family of proteins. Lining the inner surface of the inner nuclear membrane, lamin A/C are a major component of the nuclear lamina, a proteinaceous meshwork that performs many important molecular functions^1^. One such role of the nuclear lamina is to provide physical reinforcement to the nucleus, protecting its structural integrity during cellular processes that mechanically stress the nucleus, as well as in tissues subjected to constant mechanical duress. Lamin A/C are encoded by the *LMNA* gene, for which lamin A and C are produced from a single *LMNA* transcript through alternative splicing of the preRNA^2^. A notable aspect of the *LMNA* gene is that despite ubiquitous expression in most differentiated mammalian somatic cells, mutations incurred cause a spectrum of human diseases loosely grouped by the affected tissue termed laminopathies^3^. The most frequently reported laminopathy in patients is cardiomyopathy (herein referred to as *LMNA* cardiomyopathy), characterized by variable age of onset and severity^4-7^. Similar variability in the onset and severity of pathological fibrosis is also observed and is in many cases absent^4,7-14^, indicating that either modifier genes exist or unidentified factors contribute to the disease pathogenesis and/or progression. This incomplete understanding of the pathogenic mechanism needs to be addressed for translation of therapies aiming to preserve cardiac muscle.

One of the unmet challenges to gain meaningful insights into the mechanisms underlying *LMNA* cardiomyopathy is to elucidate how various cell types of the heart are affected by *LMNA* mutations and, more importantly, how these cell type-specific pathogenic effects are integrated at the tissue level to engender cardiomyopathy. In addition to cardiomyocytes (CMs), the heart is comprised of various stromal cells such as cardiac fibroblasts (CFs), endothelial cells, smooth muscle cells, neurons, pericytes, adipocytes, and resident/infiltrating leukocytes that all express lamin A/C^15^. Therefore, they can all uniquely contribute to the *LMNA* cardiomyopathy pathogenesis and/or progression. Integrative analyses from cell type-specific approaches would assist in building a foundation upon which successful treatments can be devised.

To this end, several cell type-specific studies targeting lamin A/C expression in CMs have been recently reported^16-19^ including our own^20^. Despite using different strategies to deplete lamin A/C in perinatal^16,19^ and adult^17,18,20^ CMs, there is a remarkable consistency in the presentation of the disease phenotype; severe deterioration of cardiac function accompanied by robust pathological fibrosis developed by 4 weeks following CM-specific lamin A/C depletion. In our own study, using a novel Cre driver mouse line capable of translatome profiling, we observed significant damage to the CM nuclear envelope (NE) and the Golgi, which preceded the decline in cardiac function and robust pathological fibrosis^20^. These results suggest that the aggressive fibrosis development observed in the CM-specific deletion model may be enabled by the intact expression of wildtype (WT) lamin A/C in CFs. Moreover, the extent of the NE and the Golgi damage arising from *Lmna* deletion may directly disrupt CF function, which may indirectly influence the rate of disease progression and/or severity.

To begin to explore the functional relevance of lamin A/C in CFs, we employed both *in vivo* and *in vitro* models to elucidate whether CF functions are compromised by *Lmna* mutations. In response to myocardial damage, normally quiescent CFs are activated to proliferate, migrate to the site of injury, and upregulate the expression of extracellular matrix and matricellular proteins. However, studying primary CFs poses a technical challenge as cellular states of CFs are easily perturbed, especially under *in vitro* culture on stiff surfaces^21,22^. Under these conditions, CFs spontaneously differentiate into myofibroblasts^21,22^, which are activated CFs that have acquired contractile machinery by expressing α-smooth muscle actin^23^ (αSMA). We developed a novel approach using defined factors to maintain CFs in their inactivated state despite prolonged culture in tissue culture plasticware. By leveraging this method, we elucidate whether the aforementioned functions of CFs are compromised (perhaps even enhanced) by *Lmna* mutations. Furthermore, we intercrossed our CM-specific mouse model^20^ with two different fibroblast-selective Cre recombinase driver lines^24,25^ to simultaneously deplete lamin A/C in CMs and CFs in the heart. Our studies using these *in vivo* and *in vitro* models indicate a counterbalance between CFs and CMs in driving the rate of progression and the severity of *LMNA* cardiomyopathy.

## Results

### Suppression of spontaneous activation of CFs using defined supplements during in vitro culture

To study the function of lamin A/C in CFs *in vitro*, we began by initiating the culture of primary adult human (hCFs) and murine CFs (mCFs) isolated from 12-week-old *Lmna*^flox/flox^ mice with the eventual goal of *in vitro* knockdown of lamin A/C expression. Both hCFs and mCFs were vimentin positive, maintained spindly morphology typical of non-activated CFs (Supplemental Fig. 1A), and expressed relatively high levels of *TCF21* and *COL1A1* (Supplemental Fig. 1B). In response to 10 ng/ml transforming growth factor (TGF)-β, hCFs assumed a larger and more flattened morphology with stress fibers comprised of αSMA (Supplemental Fig. 1C), which are features consistent with myofibroblast differentiation. Similar results were obtained with mCFs (data not shown).

CFs spontaneously differentiate into myofibroblasts under *in vitro* culture on hard plasticware, largely by the autocrine activation of TGF-β coupled with mechanotransduction-mediated MRTF-SRF signaling^26-28^ (Fig. 1A). The current benchmark for the determination of the CF differentiation states (in addition to cell morphology, nuclear size, and gene expression signatures) is the emergence of two different actin isoforms in sequential order. Resting CFs differentiate into proto-myofibroblasts, characterized by cytoplasmic actin stress fibers, and then into myofibroblasts, which coincides with the emergence of stress fibers composed of αSMA^29,30^. We explored ways to limit the spontaneous myofibroblast differentiation of CFs grown on tissue culture plastics. A study from Dreisen and colleagues demonstrated that culturing adult rat CFs in flat tissue culture plastics with standard media (DMEM with 10% FBS) supplemented with 10 μM of y-27632 (Rho-associated coiled coil forming protein serine/threonine kinase [ROCK] inhibitor) and 3 μM of SD-208 (TGF-β receptor I inhibitor) reduced the expression of cytoplasmic and smooth muscle actin isoforms and enhanced proliferation^31^. Building on this foundation, we repeated these experiments with hCFs and mCFs and found that although these supplements in the culture media reduced the expression of αSMA, the proliferative capacity remained low (data not shown). To overcome this issue, we also included 10 ng/ml of basic fibroblast growth factor-2 (bFGF-2). We reasoned that bFGF-2 would provide dual benefits; provide a strong mitogenic signal to facilitate CF proliferation while simultaneously antagonizing TGF-β-mediated type I collagen and αSMA expression^32-38^. Indeed, the addition of these supplements (y-27632, SD-208, and bFGF2) noticeably reduced cytoplasmic actin stress fibers as well those comprised of αSMA in human (Fig. 1B, C, and D [right panel]) and mouse (Fig. 1D [left] and Supplemental Fig. 1D) CFs. Moreover, the expression of multiple genes associated with myofibroblast differentiation were all significantly inhibited in mCFs (Fig. 1E).

**Fig. 1.**
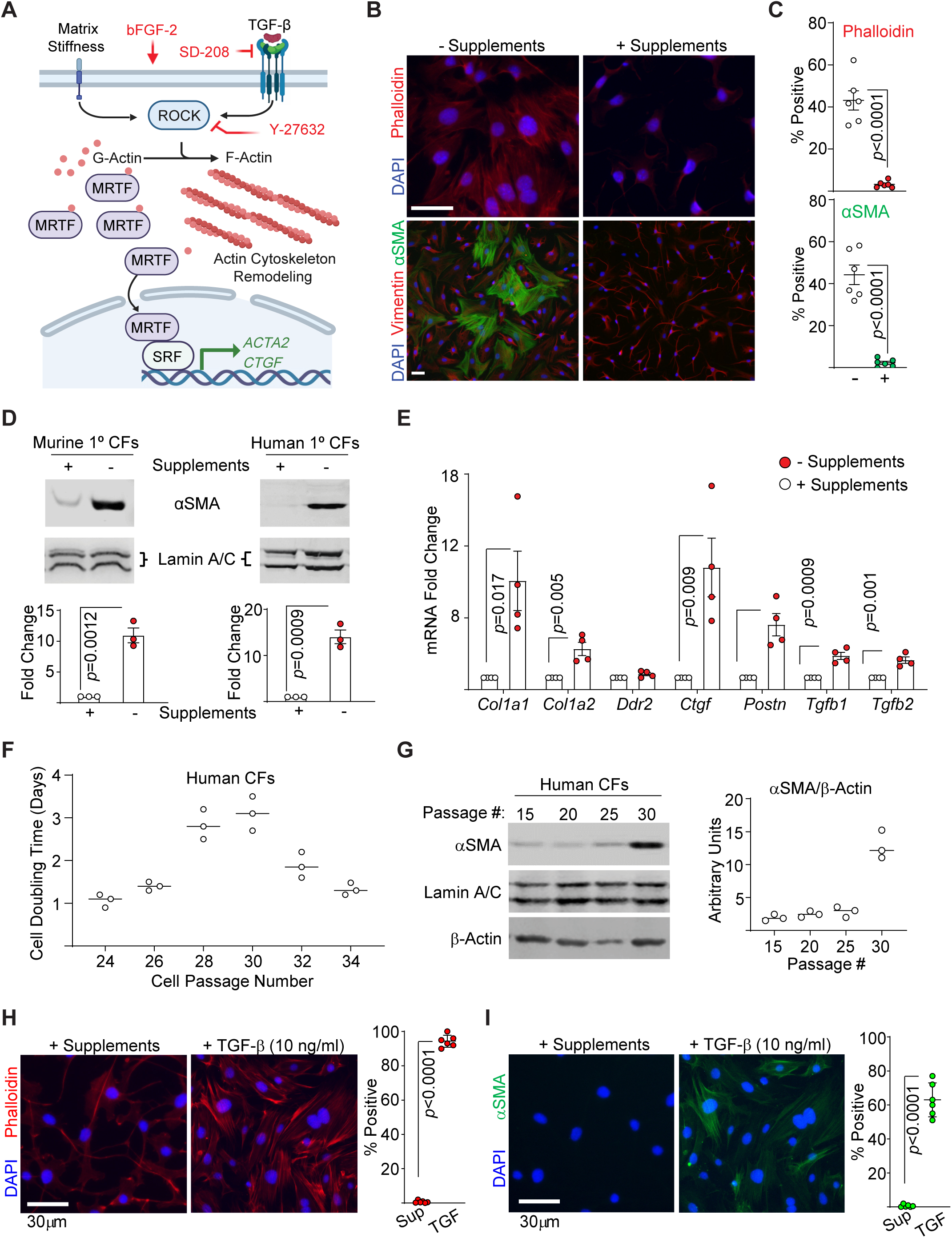
Suppression of stiffness-induced cardiac fibroblast activation using defined factors during prolonged *in vitro* culture. **A**) Schematic of stiffness-induced activation of CFs and the effects of supplements bFGF2, y-27632 (ROCK inhibitor), and SD-208 (TGF-βRI inhibitor). **B**) Immunofluorescence images of hCFs stained with phalloidin (top), and α-smooth muscle actin (αSMA) and vimentin on the bottom. DAPI was stained to visualize the nucleus. Representative images from n = 3 experiments. Scale bars = 30μm. **C**) Quantitation of phalloidin and αSMA positive cells. 6 random images from 3 experiments were analyzed as described in Methods. *p* values = unpaired, 2-tailed Student’s t-test. Error bars = SEM. **D**) αSMA and lamin A/C (loading control) immunoblots on cell extracts from mouse (left) and human (right) CFs with or without supplements. Representative blots are shown from 3 experiments. Quantitation with *p* values (unpaired, 2-tailed, Student’s t-test) are shown below the blots. Error bars = SEM relative to + supplements set to “1”. n = 3. **E**) qPCR of profibrotic markers in mRNA isolated from mCFs cultured in the presence or absence of supplements. *p* values = unpaired, 2-tailed, Student’s t-test. Error bars = SEM relative to + supplements set to “1”. n = 5. **F**) Cell number doubling time of hCFs during prolonged *in vitro* subculture from passage number 24 to 34. n = 3 experiments. **G**) Representative αSMA and lamin A/C (loading control) immunoblots on hCF extracts at different passage number cultured with supplements from n = 3 experiments. Quantitation is shown on the right. **H, I**) Immunofluorescence of hCFs stained with phalloidin (**H**), αSMA antibodies (**I**) with DAPI counterstain at passage 45 cultured with supplements or with 3-day 10 ng/ml TGF-β treatment. Representative images from n = 3 experiments. Quantitation, performed as described in 1C, is shown on the right. *p* values = unpaired, 2-tailed, Student’s t-test. Error bars = SD. n = 3.

To test the long-term effectiveness of our culture method, we maintained hCFs in long term *in vitro* subculture. At passage ∼25, despite replenishment with fresh media + supplements every two days, we noticed that the cells began to increase in size and assumed a more flattened and outspread morphology reminiscent of fibroblasts undergoing senescence (data not shown). This change in morphology was accompanied by increased cell doubling time (Fig. 1F) and expression of αSMA (Fig. 1G) despite the addition of supplements. We reasoned that perhaps we can induce spontaneous immortalization by maintaining continuous subculture^39,40^. By passage 34, we observed an outgrowth of cells that were morphologically similar to the primary CFs they were derived from (data not shown). At passage 45, immortalized hCFs expressed little to no cytoplasmic actin stress fibers and no αSMA stress fibers (Fig. 1H, I). Importantly, when stimulated with TGF-β, stress fibers composed of cytoplasmic actin and αSMA were readily observable (Fig. 1H, I), indicating that these cells retained the capacity to activate and differentiate into myofibroblasts. Hence, culturing primary adult hCFs and mCFs using our defined supplements maintains high proliferative capacity while simultaneously inhibiting cytoplasmic and αSMA stress fiber formation, as well as gene expression pattern emblematic of myofibroblasts, even during extended *in vitro* culture.

### Lamin A/C depletion disrupts various functions of CFs

We then assessed the impact of *LMNA* mutations on CF function. We focused on two differentially targeted *Lmna* mutation models; virus-mediated lamin A/C depletion (short hairpin RNA [sh1 and sh2] in hCFs and adenovirus-mediated Cre [AdCre] delivery in mCFs) and those isolated from *Lmna*^H222P/H222P^ mice, which represent a non-virally modified model with a missense mutation documented in patients^41^. The virus-mediated depletion in hCFs and mCFs achieved ∼85% (for both sh1 and sh2) and 90% knockdown, respectively (Supplemental Fig. 2A). Phenotypic assessment of CFs transiently transected with GFP-Cre expression vectors revealed irregular cell morphology with cytoplasmic protrusions (Supplemental Fig. 2B). Irregularly shaped cells were also observed in hCFs soon after infection with lentiviruses (between day 3 to day 8 post infection) delivering two different shRNA sequences (sh1 and sh2) (data not shown). We observed significant decreases in hCF proliferation following lentiviral infection with sh1 and sh2 as well as with the scrambled (ctrl) sequence, indicating that the infection itself leads to cell cycle arrest, most likely due to the activation of cellular antiviral responses (Fig. 2A). Nonetheless, the proliferation of hCFs expressing scrambled sequence recovered earlier than those expressing sh1 and sh2, suggesting that *LMNA* knockdown contributes to delayed cell proliferation. We then assessed lamin A/C levels at day 6 and day 15 post infection (Fig. 2C). At day 6, we observed a significant reduction in lamin A/C levels but this reduction was lost by day 15. These data suggest that cells with the highest lamin A/C knockdown are eventually outgrown and overtaken by those with lower knockdown levels during continuous subculture, indicating that WT lamin A/C expression is necessary for CF proliferation and/or viability. To remove potential confounding effects caused by viral infection, and to ascertain whether a point mutation causes a similar delay in cell growth, we assessed mCFs isolated from *Lmna*^H222P/H222P^ mice. mCFs isolated from *Lmna*^+/+^ and *Lmna*^H222P/H222P^ mice were adapted to tissue culture (passage 0) at which cell growth was measured for 5 days (Fig. 2C). Similar to the depletion model, mCFs carrying the H222P mutant variant displayed delayed growth relative to cells with WT lamin A/C (Fig. 2C), indicating that lamin A/C may be an important contributor to cell growth/viability in primary CFs.

**Fig. 2.**
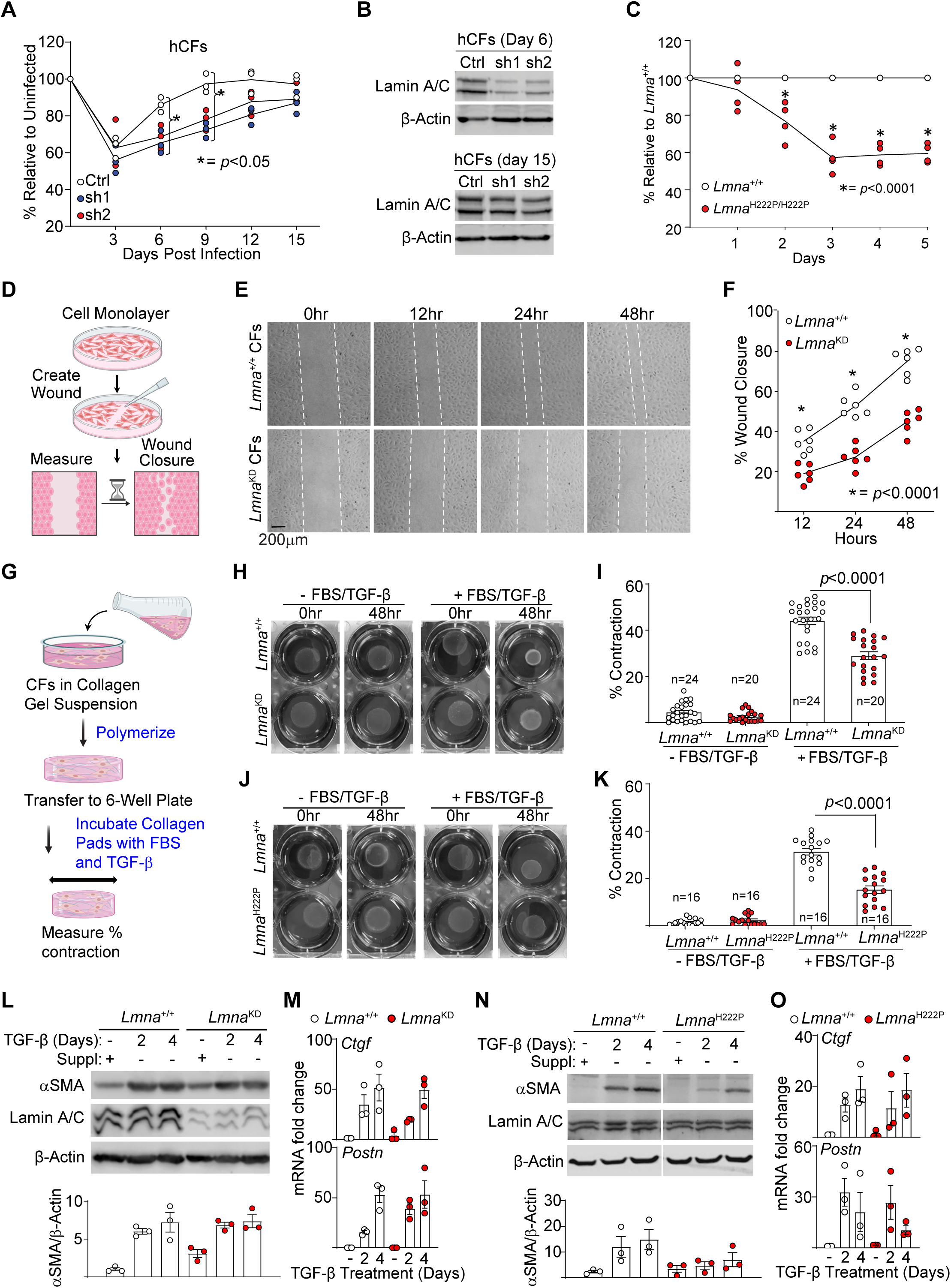
Lamin A/C mutations impair CF function. **A**) Growth kinetics of hCFs infected with lentivirus carrying blank vector (Ctrl) and two discrete short hairpin RNAs (sh1, sh2) targeting *LMNA.* The cell number was assessed in triplicates from n = 3 experiments and the data are presented as % cell number relative to controls. *p* values = two-way ANOVA with Tukey correction. **B**) Lamin A/C immunoblots on hCFs at 6 and 15 days post infection with lentiviruses carrying sh1 and sh2. β-actin was used as loading control. A representative Fig. is shown from 3 experiments. **C**) Growth kinetics of mCFs from *Lmna*^H222P/H222P^ mouse. Data presented as % cell number relative to *Lmna*^+/+^ mCFs set to 100%. *p* values = two-way ANOVA with Tukey correction. **D**) Scratch migration assay schematic, **E**) representative migration micrographs, and **F**) assay results from *Lmna*^+/+^ and *Lmna*^fl/fl^ (*Lmna*^KD^) CFs infected with AdCre presented as % wound closure relative to 0hr time point. *p* values = two-way ANOVA with Tukey correction based on n = 3 experiments performed in duplicates. **G**) Collagen pad contraction assay schematic, **H**) representative pad images, and **I**) assay results from *Lmna*^+/+^ and *Lmna*^KD^ CFs presented as % contraction relative to their respective areas at 0 hr from n = 6 experiments. 10% FBS and 10 ng/ml TGF-β was used to induce contraction. *p* value = one-way ANOVA with Tukey correction. Error bars = SEM. **J**) and **K**) show representative pad images and assay results from *Lmna*^+/+^ and *Lmna*^H222P/H222P^ CFs, respectively, with same statistical analysis as in 2I from n = 6 experiments. **L**) Representative immunoblots on extracts from *Lmna*^+/+^ and *Lmna*^KD^ mCFs probed for αSMA, lamin A/C, and β-actin (loading control) from n = 3 experiments. Bottom shows quantitation of blots. **M**) qPCR analyses on samples from 2l probed for *Ctgf* and *Postn* mRNA expression. “-”, “2”, and “4” denote untreated, 2, and 4 day 10 ng/ml TGF-β treatment, respectively. Error bars = SEM from 3 experiments. **N**) and **O**) show the same data as **L**) and **M**), respectively, but with *Lmna*^+/+^ and *Lmna*^H222P/H222P^ mCFs.

We then assessed whether *Lmna* mutations also impair CF cell migration. Prior studies using non-CF cells showed cell migration defects, largely stemming from defects in nucleo-cytoskeleton decoupling^42-45^. We presume similar defects would be evident in our primary CF models and we tested this hypothesis using a scratch migration assay (Fig. 2D-F). Interestingly, CF migration (in 2-dimensional space) appears to be impacted differentially depending on the *Lmna* mutation. For example, lamin A/C-depletion impaired cell migration in both mouse (Fig. 2E) and human (Supplemental Fig. 2C, D) CFs. However, mCFs with the H222P mutation in *Lmna* resulted in faster migration relative to WT controls (Supplemental Fig. 2E, F), suggesting that for 2-dimensional migration, H222P lamin A/C may confer a gain-of-function advantage.

### Lamin A/C depletion and H222P mutation impair CF contractile ability

We next examined CF contraction (Fig. 2G-K and Supplemental Fig. 2G, H). In response to myocardial damage, activated CFs attain contractile ability by expressing αSMA^23^. We tested whether CF contraction is altered in response to *Lmna* mutation by performing collagen pad contraction assays. *Lmna*-deleted mCFs displayed significantly reduced collagen pad contraction relative to their WT counterparts (Fig. 2H, 2I). Similarly, hCFs depleted of lamin A/C displayed similar contractile impairments (Supplemental Fig. 2G, H). We repeated the same analysis on mCFs isolated from *Lmna*^H222P/H222P^ mice to assess the impact of H222P mutant variant expression. As was the case for the depletion models, the H222P mutation also impaired CF contraction (Fig. 2J, K). These results indicate that despite differences in *Lmna* mutations, H222P mutation behaves similarly to the loss-of-function depletion models for contraction.

### TGF-β signaling is largely intact in CFs harboring Lmna mutations

As TGF-β plays a pivotal role in CF activation and differentiation into myofibroblasts, we examined the protein expression of αSMA as well as the genes encoding matricellular proteins *Ctgf* and *Postn* in response to 10 ng/ml TGF-β. At 2 and 4 days TGF-β stimulation, lamin A/C-depleted mCFs (Fig. 2L, N) and hCFs (Supplemental Fig. 3A) exhibited comparable αSMA protein expression relative to the WT counterparts. The induced mRNA expression of *Ctgf* and *Postn* were also comparable between WT and lamin A/C knockdown CFs (Fig. 2M, O, and Supplemental Fig. 3B, C). Similarly, TGF-β signaling responses, as determined by phosphorylated Smad2 and Smad3 (pSmad2/3), were also comparable (Supplemental Fig. 3D). Reducing the TGF-β concentration (100 pg – 1 ng/μl) had little to no effect on the relative responses between WT and *Lmna* mutant CFs (data not shown).

In contrast, mCFs expressing the H222P mutant variant of lamin A/C displayed slightly reduced αSMA protein and *Postn* mRNA expression, while *Ctgf* mRNA levels, albeit variable, were not significantly different than WT CFs (Fig. 2N, O; contiguous blot images are shown in Supplemental Fig. 3E). Our subcellular fractionation studies to detect pSmad2/3 in the nuclear and chromatin bound fractions revealed that, although pSmad2/3 enter the nucleus at comparable levels, they appear to be released from the chromatin fraction earlier in *Lmna*^H222P/H222P^ mCFs than in the WT counterparts (Supplemental Fig. 3F), which may underlie the slightly dampened TGF-β response. Altogether, these studies demonstrate that CF functional defects arising from *Lmna* mutations depend on both the type of *Lmna* mutation as well as the cellular function being tested.

### Golgi abnormalities associated with CFs with Lmna mutations

We previously showed that lamin A/C depletion in adult CMs causes perinuclear damage affecting the Golgi^20^. Despite the significant morphological and structural differences between these two cell types, we reasoned that perhaps similar perinuclear damage may be present in CFs harboring *Lmna* mutations. To test our hypothesis, we assessed the correlation between the Golgi size and severity of the nuclear damage in AdCre-infected *Lmna*^+/+^ and *Lmna*^fl/fl^ CFs cultured on glass chambered slides. The Golgi size, determined by the area of Golgi-resident protein RCAS1 as a surrogate Golgi marker, were plotted against nuclear damage severity, determined and expressed as nuclear irregularity index (NII) using nuclear morphometric analysis^46^, where higher NII values denote more irregular nuclei. We observed a more negative correlation between the Golgi footprint and NII in CFs with AdCre infected *Lmna*^fl/fl^ (Pearson r = -0.47) relative to *Lmna*^+/+^ CFs (Pearson r = -0.17) (Fig. 3A, B). This indicates that lamin A/C-depleted CFs with the most severe nuclear damage also exhibit a smaller Golgi footprint. Consistent with our findings in CMs^20^, these results indicate that the severity of the nuclear damage also reduce the Golgi.

**Fig. 3.**
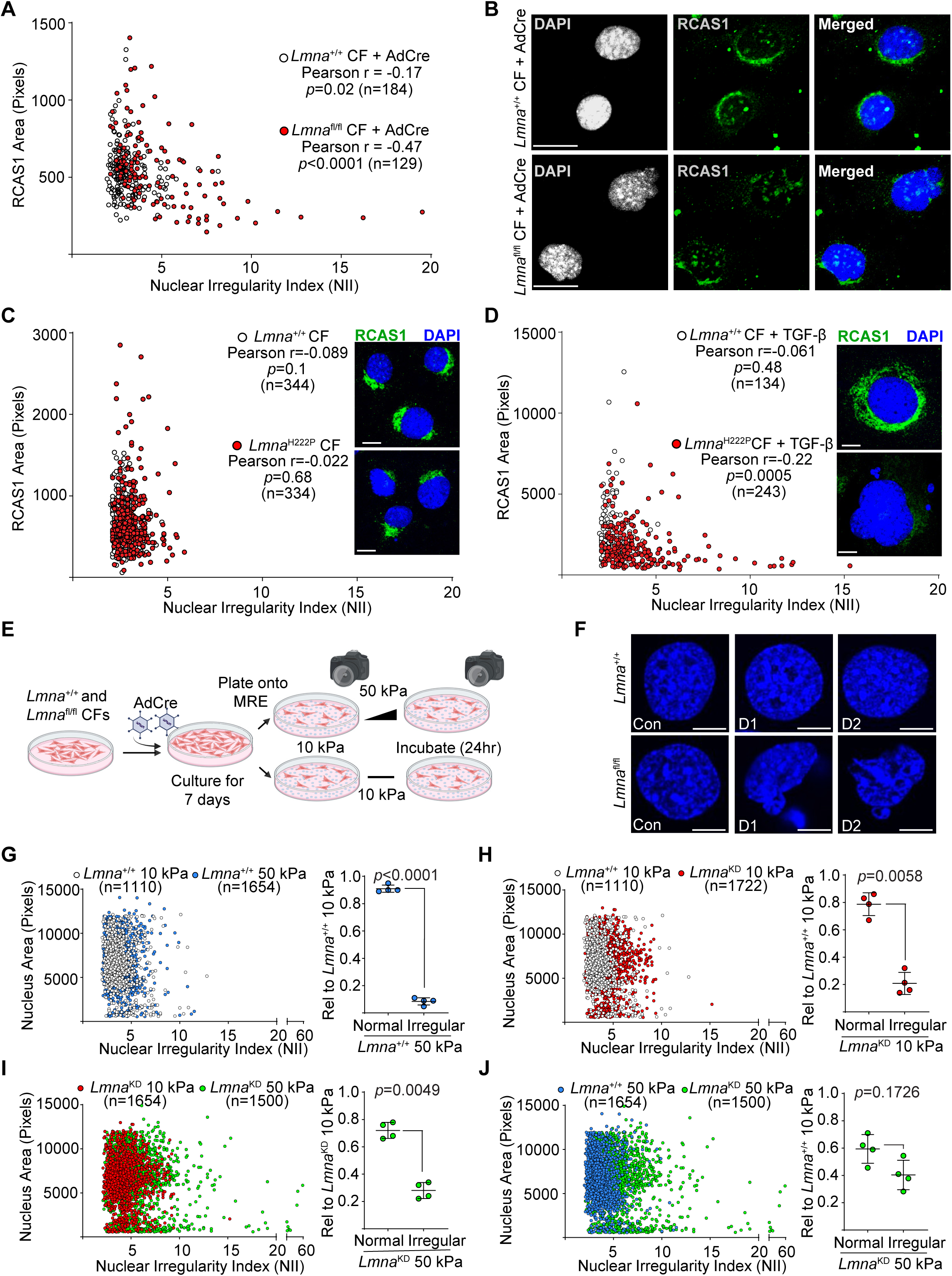
*Lmna* mutations underlie perinuclear damage in CFs. **A**) NII plotted against perinuclear RCAS1 signal area from AdCre-infected *Lmna*^+/+^ and *Lmna*^fl/fl^ mCFs. Data points were compiled from 3 experiments. Pearson r *p* values (two tailed) are shown. **B**) Representative micrographs of DAPI and RCAS1 stained mCFs from 3A. Scale bars = 20 μm. **C, D**) NII plotted against perinuclear RCAS1 signal area from *Lmna*^+/+^ and *Lmna*^H222P/H222P^ mCFs (**C**) and *Lmna*^+/+^ and *Lmna*^H222P/H222P^ mCFs stimulated with 10 ng/ml TGF-β for 48 hr (**D**). Representative micrographs of DAPI and RCAS1 stained mCFs are shown in line with the plotted graph. Scale bars = 20 μm. **E**) MRE study design schematic exploiting contact inhibition. **F**) Representative nuclear structure of AdCre-infected *Lmna*^+/+^ and *Lmna*^fl/fl^ mCFs under contact inhibition (Con) as well as day 1 (D1) and day 2 (D2) after replating to allow cell migration/replication. Scale bars = 10 μm. **G-J**) NII plotted against nuclear size after 24 hr culture in MRE with **G**) AdCre-infected *Lmna*^+/+^ at 10 and 50 kPa; **H**) AdCre-infected *Lmna*^+/+^ and *Lmna*^fl/fl^ at 10 kPa; **I**) AdCre-infected *Lmna*^fl/fl^ at 10 and 50 kPa, and **J**) AdCre-infected *Lmna*^+/+^ and *Lmna*^fl/fl^ at 50 kPa. The data points were compiled from n = 4 experiments, which are reflected in the quantification of normal and irregular nuclei (described in more detail in Methods) shown on the right of each plot. The decreasing relative differences (as denoted by increasing *p* values) demonstrate a relative shift from normal to irregular nuclei between the comparisons. *p* values = paired, 2-tailed, Student’s t-test. Error bars = SD from 4 independent experiments.

We carried out similar analysis on *Lmna*^H222P/H222P^ CFs, which express the point mutant lamin A/C at comparable levels as the WT controls. Under basal conditions (with additives), the Golgi in *Lmna*^H222P/H222P^ CFs appeared mostly normal, although the presence of abnormally shaped Golgi (deviation from the distinct crescent shape) was noticeable (Fig. 3C micrographs). Furthermore, there was no correlation between the reduced Golgi footprint and increased NII (Fig. 3C). We then performed the analysis after 48 hr TGF-β (10 ng/ml) stimulation, which stimulates CF differentiation into myofibroblasts. Following TGF-β stimulation, we noted two obvious changes in a significant portion of the cells; both the nucleus (Supplemental Fig. 4A, B) and the Golgi (Supplemental Fig. 4C) became enlarged, likely to accommodate the acquisition of cellular functions typical of myofibroblasts such as the robust production and release of matricellular and extracellular matrix proteins. Notably, the negative correlation between the Golgi size and NII was only observed in *Lmna*^H222P/H222P^ CFs with TGF-β treatment (Fig. 3D). This indicates the forces exerted on the nucleus during myofibroblast differentiation are needed to unmask the structural abnormalities of the nuclear lamina composed of the H222P variant of lamin A/C. In contrast, the detrimental impact of lamin A/C depletion on the NE integrity is relatively greater in severity, such that these perturbations are evident even at the basal condition. To validate our results with RCAS1, we repeated our analyses using antibodies against an alternate Golgi marker 58K^47^. Similar results were observed, both in the KD (Supplemental Fig. 4D, E) and the H222P mutant models (Supplemental Fig. 4F, G), with and without TGF-β stimulation. Our collective data indicate that although lamin A/C depletion in CFs elicits similar perinuclear damage as observed in CMs, the penetrance of *Lmna*^H222P/H222P^ mutation on the nuclear integrity requires TGF-β stimulation.

### Pathophysiological matrix stiffness induces nuclear abnormalities in CFs with Lmna mutations

One of the technical hurdles we encountered studying lamin A/C knockdown models using AdCre in standard tissue culture plasticware is the stochastic nature of the knockdown, causing asynchronous depletion of lamin A/C in a given population of cells. Although ∼3 days post AdCre infection are typically required to achieve lamin A/C KD (Supplemental Fig. 5A), asynchronous depletion is evident at 48hr (Supplemental Fig. 5B). This causes corresponding asynchronous nuclear damage, confounding accurate assessment of the impact of lamin A/C loss on nuclear damage. Prior studies indicated that cellular processes such as proliferation and migration generate forces that can elicit nuclear damage, particularly in lamin A/C silenced cells^48-50^. Given that our CFs are primary in nature, these cells still retain contact inhibition. We leveraged this insight to synchronize the nuclear damage by infecting *Lmna*^fl/fl^ mCFs with AdCre at ∼70% confluency, such that, by the time lamin A/C depletion is achieved, the CFs reach 100% confluency and enter contact-inhibited state. The inability to migrate or proliferate should limit nuclear damage after lamin A/C knockdown until the cells are replated, allowing synchronization of nuclear damage.

To test the effectiveness of our strategy, we utilized stiffness-tunable matrix constructed from magnetorheological elastomers (MRE). These MRE substrates are constructed from polydimethylsiloxane (PDMS) infused with iron particles that can be dynamically tuned to physiologically relevant stiffness from ∼10 kPa (normal cardiac tissue) to 50 kPa (fibrotic cardiac tissue) by varying the magnetic field using an external magnet^51^. *Lmna*^+/+^ and *Lmna*^fl/fl^ CFs, infected with AdCre at ∼70% confluency, were allowed to recover and grow to full confluency for 5 days (Fig. 3E). They were then replated onto MRE substrates maintained at 10 kPa and allowed to attach overnight. The substrate stiffness was then raised to 50 kPa using an external magnet while the control plate was maintained at 10 kPa. After 24 hr, Hoechst 33342-stained nuclei from were imaged and analyzed to determine NII. Prior to experiments with MREs, we tested our strategy by replating Adcre-infected *Lmna*^+/+^ and *Lmna*^fl/fl^ (*Lmna*^KD^) CFs onto regular plasticware and assessing the emergence of nuclear damage (Fig. 3F). As expected, while both *Lmna*^+/+^ and *Lmna*^KD^ CFs retained normal nuclear structure during contact inhibition, nuclear damage emerged only in the *Lmna*^KD^ CFs 24 hr after replating onto plastic cultureware (Fig. 3F). This is further validated by our results using the MREs; greater than 90% of *Lmna*^+/+^ CFs retained normal nuclear shape when the MRE stiffness was raised from 10 to 50 kPa (Fig. 3G), confirming that CFs with WT lamin A/C expression are resistant to nuclear damage. This percentage decreases to ∼80% when comparing *Lmna*^+/+^ CFs at 10 kPa relative to *Lmna*^KD^ CFs at the same stiffness (Fig. 3H), indicating that although overt nuclear damage in *Lmna*^KD^ CFs is largely inhibited at 10 kPa, some damage nonetheless still occurs at this substrate stiffness. Despite the increased baseline damage in *Lmna*^KD^ CFs at 10 kPa, further damage to the nucleus was observed at 50 kPa (Fig. 3I; ∼28% irregular nuclei relative to 10 kPa). Notably, when compared to *Lmna*^+/+^ CFs at 50 kPa, *Lmna*^KD^ CFs at the same stiffness represents over 40% increase in nuclear damage (Fig. 3J). Taken together, our results demonstrate that 1) nuclear damage from lamin A/C depletion occurs in pathophysiologically relevant stiffness and 2) our strategy using contact inhibition to synchronize nuclear damage is viable.

### Concurrent deletion of Lmna in CMs and CFs achieves salutary effects on the heart

We and others recently reported that *Lmna* deletion in CMs *in vivo* resulted in rapid yet consistent development pathological fibrosis by 4 weeks post tamoxifen (Tam), regardless of the developmental stage of the CMs^16,17,19,20^. Given this remarkable consistency in the kinetics of the fibrotic response, coupled with our *in vitro* studies showing disruption of CF function with *Lmna* deletion, we reasoned that lamin A/C in CFs may be an important determinant of the rate and the severity of *LMNA* cardiomyopathy development mediated through fibrosis. To determine the relevance of *Lmna* in CFs specifically during *LMNA* cardiomyopathy pathogenesis, we interbred our CM-only *Lmna* deletion mice (CM-CreTRAP:*Lmna*^fl/fl^) with two CF-selective Cre drivers; *Col1a2*-CreERT^25^ referred to as CF(C)-Only and *Postn*^MCM^ mice^52^ as CF(P)-Only, the latter of which would enable *Lmna* co-deletion specifically in activated CFs (Fig. 4A). This approach effectively generates bigenic Cre driver lines (CM-CreTRAP:*Col1a2*-CreERT:*Lmna*^fl/fl^ [herein referred to as CM:CF(C)] and CM-CreTRAP:*Postn*^MCM^:*Lmna*^fl/fl^ [CM:CF(P)] with Tam-regulatable Cre to delete *Lmna*^fl/fl^ in both CMs and CFs. These mice were then subjected to Tam dosing at 12 weeks of age according to the treatment strategy outlined in Fig. 4A. Treatment groups with mice carrying the *Postn*^MCM^ allele were also put on Tam chow to ensure the presence of Tam when the endogenous *Postn* promoter is robustly activated at ∼2 weeks post initial Tam injection^20^ (Fig. 4A). At 2 and 4 weeks post Tam, we performed M-mode echocardiography (echo) to assess cardiac performance by measuring systolic and diastolic diameters of the left ventricle, which were used to calculate fractional shortening (% reduction of left ventricular dimension at systole) and ejection fraction (estimated calculation of % of blood ejected from left ventricle).

**Fig. 4.**
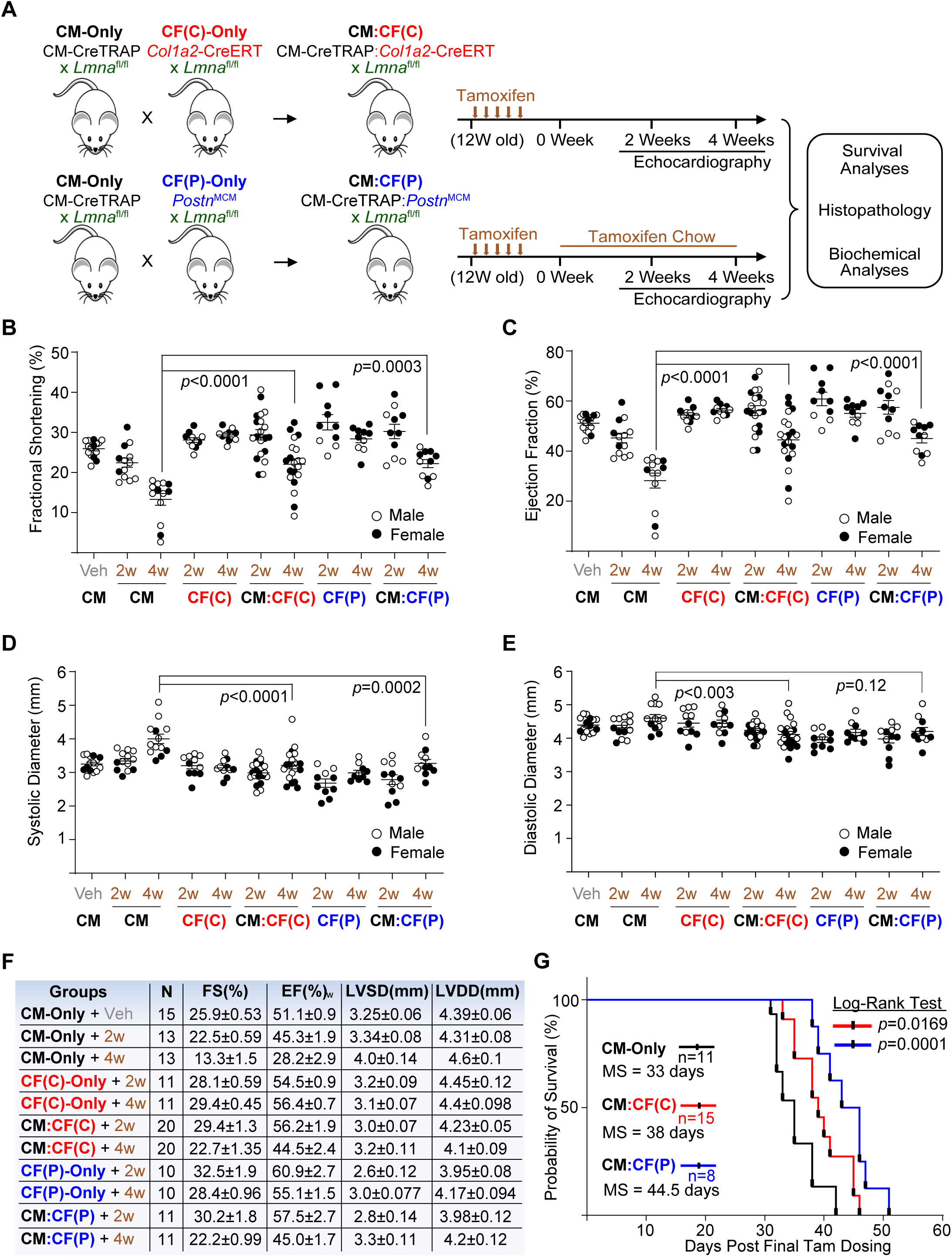
Simultaneous deletion of *Lmna* in CMs and CFs delays cardiomyopathy and enhances survival. **A**) Schematic of *in vivo* study design using intercross of CM-Only model with CF-selective, Tam-regulatable Cre driver lines. The initial administration of tamoxifen was delivered intraperitoneally (IP) at 100 mg/kg for 5 consecutive days when mice were 12 weeks old. **B-E**) Echo analysis showing fractional shortening (**B**), ejection fraction (**C**), systolic diameter (**D**), and diastolic diameter (**E**) of the left ventricle. “Veh” denotes CM-Only line at 4 weeks post corn-oil treatment delivered IP. “2w” and “4w” denote 2 and 4 weeks post IP Tam dosing. *p* values = one-way ANOVA with Tukey correction. Error bars = SEM. Compiled data from 3 independent experiments are shown. **F**) Tabulation of echo data ± SEM. FS = fractional shortening; EF = ejection fraction; LVSD = left ventricular systolic dimension; LVDD = left ventricular diastolic dimension. **G**) Kaplan-Meier plot of CM-Only, CM:CF(C), and CM:CF(P) mice. MS = median survival. *p* values were calculated using log-rank test with the survival curve of CM-Only as the comparison control.

Consistent with our previous report^20^, we observed a reduction in fractional shortening (Fig. 4B) and ejection fraction (Fig. 4C) in CM-Only group at 4 weeks post Tam relative to vehicle-treated group. In contrast, Tam-treated CF(C) did not display cardiac dysfunction (Fig. 4B, C), despite a ∼60% reduction in lamin A/C protein levels (Supplemental Fig. 6A) in the CFs of CF(C) mice, as well as the presence of the recombined floxed *Lmna* allele in the heart (Supplemental Fig. 6B). These results suggest that lamin A/C depletion in CFs, in and of itself, does not lead to cardiac pathology and that the fibrosis development in the CM-Only mice^20^ occurs as a reaction to CM damage. Identical results were obtained with CF(P) mice (Fig. 4B, C) although it is likely that the *Postn* promoter-driven MerCreMer expression did not activate in the CF(P)-Only setting. Nevertheless, we conclude that lamin A/C depletion in CFs exhibit no pathological phenotype and that Tam toxicity, although well established to occur, does not account for the cardiac dysfunction in the CM-Only mice.

We then assessed *Lmna* deletion simultaneously in CMs and CFs. Remarkably, we observed a preservation of both fractional shortening and ejection fraction at 4 weeks post Tam in both CM:CF(C) and CM:CF(P) relative to CM-only group (Fig. 4B, C). A closer examination revealed that this improvement was primarily mediated by the preservation of systolic dimension, leading to improved cardiac performance (Fig. 4D, E). We did not observe sex-dependent differences. A full list of tabulated numbers is shown in Fig. 4F. To determine whether the observed improvement in cardiac function extends lifespan, we performed survival analyses (Fig. 4G). Consistent with the echo data, both CM:CF(C) and CM:CF(P) mice displayed enhanced survival (Fig. 4G) although they both eventually succumbed to the disease, indicating that lamin A/C depletion in CFs delays, but not abrogates, disease progression.

### CM:CF Lmna co-deletion dampens pathological remodeling and fibrosis

To correlate the preservation of cardiac function with putative changes at the molecular level as well as in tissue structure, we performed RT-qPCR to measure profibrotic gene expression and histopathology (H&E, Masson’s trichrome). As we did not observe sex-dependent differences in cardiac function, tissues from both sexes were analyzed together in subsequent studies. We previously noted significant myocyte swelling/loss with abundant cytoplasmic vacuolation and interstitial fibrosis in hearts of CM-only *Lmna* deleted mice at 4 weeks after Tam^20^ and we observed similar findings here (Fig. 5A, B). In contrast, hearts from both CM:CF(C) and CM:CF(P) mice displayed reduced CM swelling/loss and vacuolation as well as significantly reduced collagen deposition (Fig. 5A, B, and Supplemental Fig. 6C). CF(C) and CF(P) histopathology was indistinguishable from negative controls (Supplemental Fig. 6D). Consistent with the histological analyses, mRNA expression analyses revealed significantly reduced levels of cardiac stress markers (*Nppa*, *Nppb*) as well as matricellular (*Postn*, *Ctgf*) and extracellular matrix proteins (*Col1a1*, *Col1a2*, *Col3a2*, *Fn1*-EDA) in hearts of both CM:CF *Lmna* deleted mice relative to the CM-only at 4 weeks post Tam (Fig. 5C). These results are indicative of impaired cardiac fibroblast function.

**Fig. 5.**
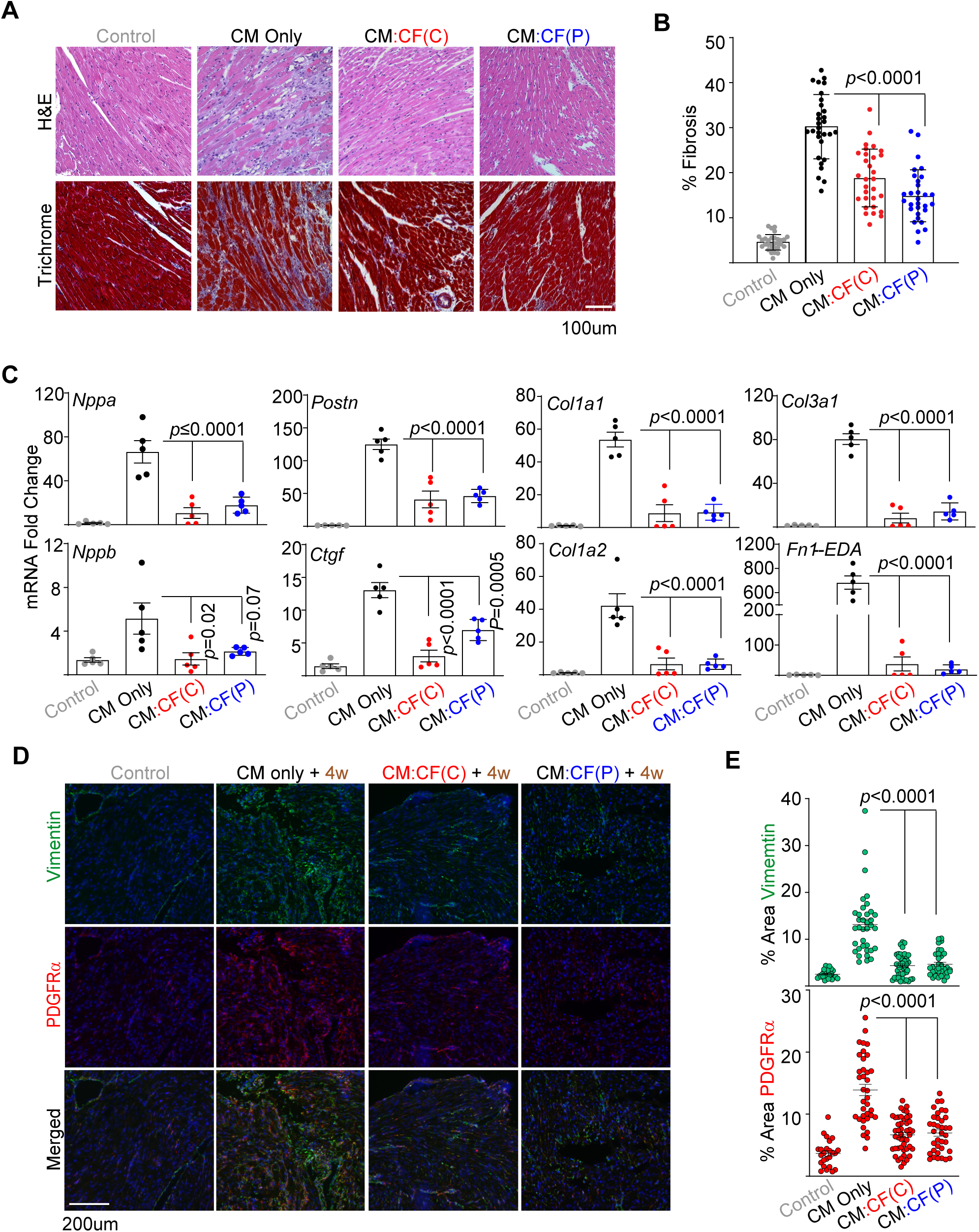
Simultaneous deletion of *Lmna* in CMs and CFs dampens fibrosis relative to CM-Only deletion. **A**) Hematoxylin and Eosin (H&E) and Masson’s trichrome staining analyses of hearts from control (CF(C)-Only), CM-Only, CM:CF(C), and CM:CF(P) mice at 4 weeks post Tam. A representative image per group (n = 3) are shown. **B**) Quantitation of % fibrosis of Masson’s trichrome staining from 30 independent images from 6 mouse hearts per group. *p* value = one-way ANOVA with Tukey correction. Error bars = SD. **C**) qPCR analyses of cardiac stress and profibrotic markers in hearts from control (CF(C)-Only), CM-Only, CM:CF(C), and CM:CF(P) mice at 4 week post Tam. Data for each group (including control) are presented as fold change relative to the mean value of the control group. *p* values = one-way ANOVA with Tukey correction. n = 5. Error bars = SEM. **D**) Representative immunofluorescence images of vimentin and PDGFRα with DAPI counterstain on hearts from control (CF(C)-Only), CM-Only, CM:CF(C), and CM:CF(P) mice at 4 weeks post Tam. **E**) Quantitation of vimentin and PDGFRα presented as % area per image field from control (n = 24 images), CM-Only (n = 36), CM:CF(C) (n = 45), and CM:CF(P) (n = 36) mice at 4 weeks post Tam from 6 mice per group. *p* values = one-way ANOVA with Tukey correction. Error bars = SEM. Data from both sexes are shown in this Fig..

Given the impact of lamin A/C depletion on CF function, we reasoned that this may lead to reduced CF content that may underlie the observed dampening of pathological remodeling and fibrosis. To assess the abundance of CFs *in situ* during *LMNA* cardiomyopathy pathogenesis, we performed immunofluorescence staining/microscopy on heart sections from Tam-treated CM:CF(C) and CM:CF(P) mice that were co-stained for vimentin and PDGFRα, both of which are CF-selective markers. In CM-only mice, we observed significant increases in CF-selective staining at 4 weeks post Tam (Fig. 5D, E). Notably, these increases were blunted in hearts of both of the CM:CF *Lmna* deletion models (Fig. 5D, E), suggesting that the CF content is reduced in the myocardium of these mice. Taken together, our results show that concurrent *Lmna* deletion in the CM:CF mice impair CF expansion to mount a robust fibrotic response.

### Distinct nuclear shape abnormalities in CM:CF Lmna co-deletion compared to CM-only deletion

During histopathological analyses, we observed an abundance of abnormal nuclei in the heart sections of CM:CF mice that were distinct from those of CM-only *Lmna* deletion. Whereas predominantly irregularly shaped and ruptured CM nuclei were noted in the CM-only hearts^20^, those from the CM:CF models exhibited oblong, but intact, nuclei (Fig. 6A), reminiscent of those observed in hearts of germline *Lmna* mutant mice^53,54^. Quantification of elongated nuclei confirmed their increased presence in the CM:CF(C) and CM:CF(P) mice but not in the CM-only mice (Fig. 6B).

**Fig. 6.**
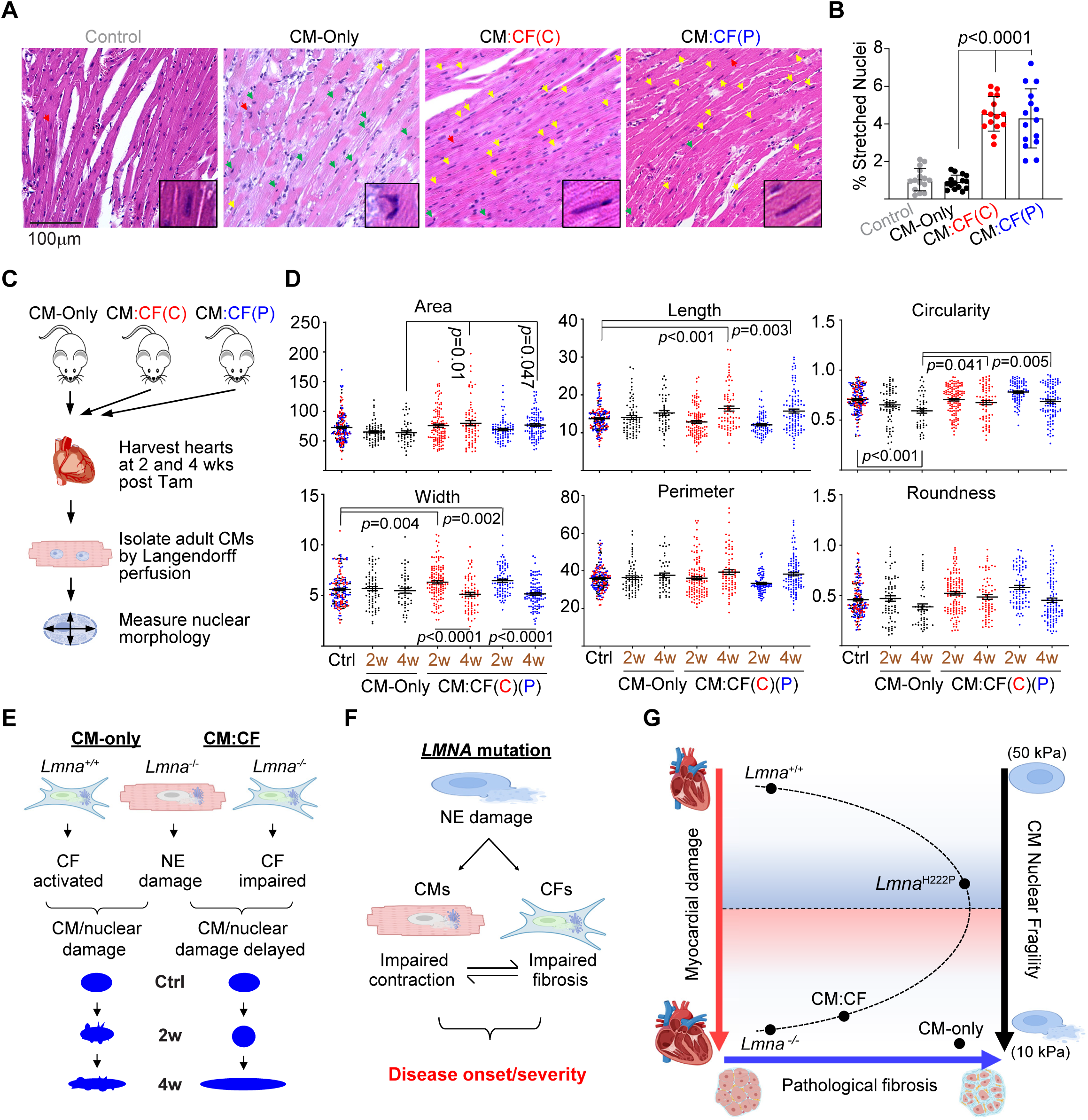
Distinct nuclear morphology associated with CM-Only relative to CM:CF models. **A**) Representative H&E images of longitudinal sections of hearts from control (CF(C)-Only), CM-Only, CM:CF(C), and CM:CF(P) mice at 4 weeks post Tam. The nucleus in the inset (red arrow) shows unique morphology typically found in each group. Green and yellow arrows denote misshapen/irregular and elongated nuclei, respectively. **B**) Quantitation of % stretched nuclei per field from 15 images from 3-5 mice per group. *p* values = one-way ANOVA with Tukey correction. Error bars = SD. **C**) Schematic of study design for CM nuclear shape analysis. **D**) CM nucleus shape measurements from control (Ctrl), CM-Only, CM:CF(C), and CM:CF(P) mice at 2 and 4 weeks post Tam. Ctrl in this case is an aggregate of data points from vehicle-treated (post 4 weeks) CM-Only and CF-Only models (both CF(C) and CF(P)) at 4 weeks post Tam. Area, width, length, and perimeter were measured in microns. Circularity and roundness index are measured from 0 to 1 with 1 denoting perfect circularity/roundness. Ctrl n = 184 CM nuclei, CM-Only 2w/4w n = 70/49, CM:CF(C) 2w/4w n = 126/74, and CM:CF(P) n = 83/102 from 3 mice per group. *p* values = one-way ANOVA with Tukey correction. Error bars = SEM. Data from both sexes are shown in this Fig.. **E**) Graphical interpretations of data from 6d. **F**) Simplified schematic of *LMNA* cardiomyopathy at the cellular level. **G**) Proposed working hypothesis of *LMNA* cardiomyopathy at the tissue level. The extent of the myocardial damage (red arrow) governed by the extent of pathological fibrosis (blue arrow) and propensity for CM nuclear damage (black arrow) in a non-linear relationship.

To determine whether distinct nuclear shape abnormalities exist specific to the CM-only vs the CM:CF models, we performed detailed nuclear shape analyses on freshly dissociated CMs from these mice at 2 and 4 weeks post Tam, similar to those reported in the *Lmna*^H222P/H222P^ mice^55^ (Fig. 6C, D). When comparing the 4 week groups exclusively, we found that nucleus area was significantly increased in CM:CF(C) and CM:CF(P) relative to CM-Only (Fig. 6D with representative images in Supplemental Fig. 6D), which is consistent with our histological observations (Fig. 6A). Moreover, we observed that circularity, which denotes local curvature of the nucleus, is significantly reduced in CM-only at 4 weeks post Tam relative to both CM:CF models (Fig. 6D). This is consistent with the increased frequency of ruptured, misshapen, and/or herniated nuclei in the 4 week CM-only group that would deviate from smooth curvature, whereas these features are much less prominent in the CM:CF models. Although the nuclear length in the 4 week CM:CF models were not significantly different from CM-only, they were significantly longer relative to controls (Fig. 6D), suggesting that the CM:CF nuclei become more elongated than those in CM-Only. Finally, the nuclear width of the CM:CF models undergo interesting transitions; at 2 weeks, the width significantly increases from controls. However, this is followed by a significant decrease at 4 weeks (Fig. 6D). This change in nuclear width was not observed in the CM-Only group. No significant differences between meaningful comparisons were observed for perimeter and roundness (which denotes aspect ratio of the nucleus). These results reveal nuclear damage processes that are clearly distinct (Fig. 6E). In the CM-Only model, the nucleus gradually loses circularity, likely due to accumulating NE rupture and/or herniation, with some elongation occurring at 4 weeks post Tam. In the CM:CF models, the nucleus appears more resistant to damage that affects the local curvature of the NE. In addition, the nucleus at 2 weeks post Tam becomes much more spherical before significantly elongating at 4 weeks (Fig. 6E). These observations are consistent with a nucleus recoiling back to a spherical shape due to loss of tension at the major axis of the nucleus. Although the exact mechanisms remain to be determined, our results indicate that CM:CF models better recapitulate the nuclear morphological changes observed in germline mouse models^53,54,56^ and are associated with a less aggressive disease.

## Discussion

In this study, we explored in CFs the functional relevance of lamin A/C. CFs are arguably the central mediators of fibrotic response to cardiac injury but despite their importance, little is known about the importance of lamin A/C in homeostatic function of CFs as well as their contribution to *LMNA* cardiomyopathy pathogenesis. Our results suggest *LMNA* mutations in CFs, and possibly in other non-myocyte populations of the heart, may play a far more substantial role in disease pathogenesis than previously appreciated. Nuclear fragility arising from mutations in *Lmna* leads to similar perinuclear damage in both CMs^18,20^ and CFs, as demonstrated in this study. However, the data presented herein indicate that CMs and CFs with *Lmna* mutations contribute to disease pathogenesis in an opposing manner; whereas lamin A/C protect CMs from perinuclear abnormalities leading to myocyte damage, they simultaneously protect CF function, thereby enabling reactive fibrosis.

We propose that *LMNA* cardiomyopathy progression and severity are conferred by the following interrelated factors: the severity of the nuclear damage, the extent of the collateral damage to the perinuclear space and the Golgi, and the intended function of the affected cell type during disease pathogenesis. In this scenario, the perinuclear damage is the main driver of pathogenesis (Fig. 6F). As both CMs and CFs are similarly affected by the perinuclear damage that perturb their respective functions in the heart, we hypothesize that the integrative impact of these damaged cells underlie disease onset/severity of *LMNA* cardiomyopathy (Fig. 6F). Although our reductionist approach focused primarily on CMs and CFs and how they function in the context of the physical component of fibrosis, we surmise that additional cell populations and non-physical factors such as inflammatory cells and paracrine-mediated mechanisms, respectively, likely contribute to the disease pathogenesis as well.

Based on the foregoing, we propose a simplified two-variable pathogenesis model in which the severity of myocardial damage from *LMNA* mutations is dependent on 1) the propensity for nuclear damage (or nuclear fragility), which is directly related to the *LMNA* mutation itself, and 2) the severity of pathological fibrosis (both the rate of its development and the extent), which can indirectly impact nuclear damage by increasing myocardial tissue stiffness (Fig. 6G). In this context, we predict that the WT *Lmna*^+/+^ and the germline *Lmna*^-/-^ mutants represent the opposite ends of the myocardial damage spectrum as well as for nuclear fragility. Furthermore, we predict that fibrosis development depends on the balance between CM damage, required for CF activation, and nuclear fragility, in a non-linear relationship. *LMNA* mutations with lower propensity for nuclear damage elicit dampened stress signals emanating from CMs (due to reduced nuclear damage) while the higher propensity may impair CF function due to the severe nuclear damage despite ample stress signals from the damaged CMs.

Pathological fibrosis has been documented in germline point mutant knock-in mouse models but their onset is much more delayed^41^ or they develop a milder form^56^. Moreover, *Lmna*^Δ8-11/Δ8-11^ mice, which were originally thought to be a knockout^57^ but in a subsequent study shown to express very low levels of truncated lamin A/C^58^, do not develop fibrosis despite the development of severe cardiomyopathy and death within 4-6 weeks after birth^54^. This is in direct contrast to the observations from CM-Only models in which the CFs that retain WT lamin A/C expression display rapid and aggressive fibrosis development^16-20^. A recent study reported that mice with a conditional *Lmna* deletion in the fibroblast lineage using *Pdgfra*-Cre developed severe cardiomyopathy and premature death by 6 weeks of age^59^, which appears to contradict our findings. However, the authors reported 25% of the CMs displayed lamin A/C depletion, which confounds the actual contribution of lamin A/C in CF to cardiomyopathy pathogenesis. Furthermore, given that *Pdgfra* promoter is activated during early embryonic development^60^, this early loss of *Lmna* may have a differing outcome relative to the adult setting. This is reminiscent of how perinatal abrogation of ∼80% of CFs leads to early lethality^61^ whereas in the adult setting, this loss was shown to be cardioprotective^62^. Nevertheless, these results suggest that lamin A/C expression is important for CFs and that loss-of-function *Lmna* mutations leading to severe nuclear damage can impair CF function.

Although our study clarifies the contribution of CFs in the pathogenesis of *LMNA* cardiomyopathy, several questions remain unanswered. For example, it remains to be determined which other cell populations of the myocardium contribute to cardiomyopathy development as mentioned previously. Even within the CF population, it is unknown whether specific CF subgroups are active in *LMNA* cardiomyopathy and how they evolve during disease progression. Recent studies in single cell transcriptomics in the heart revealed at least eight to eleven different CF subpopulations arise during various cardiac pathologies including dilated cardiomyopathy^63^, diabetic cardiomyopathy^64^, myocardial infarction^65,66^, and chronic angiotensin II infusion^67^. More relevant to *LMNA* cardiomyopathy, a recent study using a similar CM-specific *Lmna* deletional approach, but with a different Cre driver line, broadly segregated CFs into two groups^18^. Despite the differing CF subgroups found in multiple cardiac pathologies, these studies all found POSTN expressing CFs as the prominent subgroup to arise during myocardial stress. As such, although it is unclear which CF subgroups are lacking lamin A/C expression in the *Col1a2*-CreERT line, our use of the *Postn*^MCM^ line clearly targets the most prominent pathology-related CF subgroup to arise. Future studies should address these knowledge gaps.

## Methods

### Animals

Thomas Jefferson University Institutional Animal Care and Use Committee approved all animal procedures under protocol #01744. The approved procedures adhered to the NIH Guide for the Care and Use of Laboratory Animals. CM-CreTRAP transgenic mice, generated in the C57BL/6 background, have been described elsewhere^20^. *Lmna*^flox/flox68^, *Col1a2*-CreERT^25^, and *Postn*^MerCreMer24^ lines have also been previously described and obtained from The Jackson Laboratory. *Lmna*^H222P/H222P^ mice^41^, from which we derived primary CFs, were provided by Dr. Gisele Bonne. All mice in the study were backcrossed to C57BL/6J for at least 8 generations and maintained in the C57BL/6J background. Genotyping was performed by standard PCR using genomic DNA isolated from tail biopsies and primer pairs according to their initial reports (CM-CreTRAP^20^, *Lmna*^flox/flox68^, and *Lmna*^H222P/H222P^ mice^41^). Genotyping for the *Col1a2*-CreERT and *Postn*^MerCreMer^ lines were performed according to the protocol available in their respective product page at http://www.jax.org. Mice were maintained on a chow diet and housed in a disease-free barrier facility with 12/12 hr light/dark cycles. Sample size for *in vivo* analyses were determined based on initial pilot experimental results. As we did not observe sex-dependent differences in the CM-CreTRAP mice^20^, both male and female mice were used in the study. To induce Cre recombinase-mediated excision, mice were treated with Tam via two routes of administration. For intraperitoneal delivery, Tam powder (ThermoFisher, cat# J63509.ME) was dissolved in corn-oil (MilliporeSigma cat# C8267) and delivered at 100 mg/kg. For oral delivery, mice were fed Tam-laced chow (Inotiv, cat# TD.130857) ad libitum, with the approximate Tam dosing of 80 mg/kg/day, assuming 3 - 4 g of daily food intake for a 25 g mouse.

### Primary cell isolation and culture

Adult murine CM isolation (from both sexes) was performed as described^69^ with a constant pressure Langendorff perfusion system using 25 μg/ml Liberase TH (MilliporeSigma cat# LIBTH-RO). Murine CFs were derived from single cell suspensions generated from the ventricles of 12 week-old wildtype C57BL/6 and *Lmna*^fl/fl^ mice using the gentleMACS Octo dissociator (Miltenyi Biotec, cat# 130-095-937). The dissected ventricles (2 mice per genotype) were halved, rinsed in ice-cold HBSS, and placed into the gentleMACS C Tube (Miltenyi Biotec, cat# 130-093-237) containing 4.7 ml of HBSS and 300 μl Collagenase II solution (final concentration of 600 U/ml, Worthington, cat# LS004177) and 10 μl DNase I solution (final concentration of 60 U/ml, Zymo Research, cat# E1011). The C Tube containing the heart pieces were placed into Octo dissociator and ran the program “m_heart_01”. Following the program completion, the samples (still inside the C Tube) were incubated at 37°C for 30 min with intermittent resuspension by agitation every 5 min. The samples were placed back into Octo dissociator and ran the program “m_heart_02”. The single cell suspension was pelleted by brief centrifugation, resuspended with 5 ml of HBSS, and put through a 70 μm mesh cell strainer. After washing the strainer with an additional 5 ml of HBSS, the cells were centrifuged at 300xg for 10 min, resuspended in 1ml of PEB buffer (PBS pH 7.2, 2 mM EDTA, 0.5% BSA) along with 10 ml of 1X Red Blood Cell Lysis Solution (Miltenyi Biotec, cat# 130-094-183), and incubated for 2 minutes at room temperature. Following 2X wash with 1 ml PEB buffer, the cells were resuspended in DMEM with 10% FBS and plated onto a 10 cm culture plate. The media was replaced 4 hr later with DMEM + 10% FBS supplemented with 10 μM of y-27632 (Selleck Chemicals, cat# S1049), 3 μM of SD-208 (Selleck Chemicals, cat# S7624), and 1 ng/ml basic FGF-2 (Prospec, cat# cyt-557). After about 5-7 days (with media change every 2-3 days), outgrowths of CFs were visible under the microscope. Non-hypertrophic human ventricular CFs were purchased from Lonza (Lonza, cat# CC-2904).

### In vitro culture and manipulation of CFs

Human and mouse CF culture method was adapted from Driesen *et al*^31^ but with modifications. Just prior to use, the culture media (DMEM with 10% FBS) were supplemented with 10 μM of y-27632 (Selleck Chemicals, cat# S1049) and 3 μM of SD-208 (Selleck Chemicals, cat# S7624) but we also included 10 ng/ml basic FGF-2 (Prospec, cat# cyt-557). They were cultured normally on culture plates that were previously coated with 0.1% gelatin for 1 hr at 37°C followed by 2X rinse with PBS. For experiments involving the exclusion of supplements, its removal was performed in a stepwise fashion. To remove supplements, CFs were cultured overnight with 10 μM of y-27632 only. On the next day, the cells were rinsed 3X with PBS followed by the addition of DMEM with 10% FBS without any supplements. Additionally, 10 ng/ml TGF-β1 was added to DMEM with 10% FBS at this step for experiments testing CF responses to this cytokine. To deplete lamin A/C, human and mouse CFs were subjected to differing approaches. For human CFs, we employed two independent shRNAs in pLKO.1 lentiviral vector identified from human *LMNA* shRNA (MilliporeSigma cat# SHCLNG-NM_170707) with the sequences 5’-GAAGCAACTTCAGGATGAGAT (sh1) and 5’-GATGATCCCTTGCTGACTTAC (sh2). The lentiviral vectors were co-transfected into 293T cells (cultured in DMEM with 10% FBS) with the packaging vectors pCMV-dR8.2 dvpr and pCMV-VSV-G^70^. pCMV-dR8.2 dvpr and pCMV-VSV-G were gifts from Bob Weinberg (Addgene plasmid #8455; http://n2t.net/addgene: 8455; RRID: Addgene_8455 and Addgene plasmid #8454; http://n2t.net/addgene: 8454; RRID: Addgene_8454). Blank controls indicate CFs infected with lentiviruses generated from an empty vector. Virus-infected cells were selected with 2 μg/ml puromycin. To delete *Lmna* in mouse CFs *in vitro*, adenoviruses carrying mCherry (AdBlank) or mCherry-Cre (AdCre) (Vector Biolabs cat# 1767 and 1773, respectively) were used at 50 MOI.

### RNA isolation and RT-qPCR

Total RNA were isolated using Direct-zol RNA kit (Zymo Research, cat# R2053) according to the manufacturer’s instructions with a minor modification as previously described^20^. Briefly, tissues solubilized in TRIzol (Zymo Research cat# R2050-1-200) were phase-separated using chloroform and the aqueous phase containing the RNA fraction was collected and mixed with 100% molecular grade ethanol at 1:1 ratio. This mixture was further processed using the Direct-zol RNA kit according to the manufacturer’s instructions. cDNA were generated from 1 µg – 500 ng of total RNA primed with a 1:1 ratio of random hexameric primers and oligodT using RevertAid RT kit (ThermoFisher Scientific cat# K1691). qPCR was run on a QuantStudio5 thermocycler system (Life Technologies) using PowerUP SYBR-green reagent (ThermoFisher Scientific, cat# A25743). Both *Gapdh* and *Rpl13a* levels were assessed to ensure even synthesis of cDNA and those with the least min/max CT value variability was used as our internal control gene to normalize qPCR data. Fold-change values in gene expression were obtained using the ΔΔCt method ^71^ and presented as fold-change over negative controls. The complete list of qPCR primer pairs in the study are listed in the Supplemental Table 1.

### Protein extraction, immunoblot analysis

Frozen cell pellets/tissues were homogenized in chilled (4°C) radioimmunoprecipitation assay (RIPA) buffer (MilliporeSigma cat# R0278) supplemented with Pierce Protease Inhibitor cocktail (ThermoFisher Scientific cat# A32963) and 1 mM of phosphatase inhibitor sodium vanadate (MilliporeSigma cat# S6508). After sonication (Dismembrator Model F60, ThermoFisher Scientific), the protein extracts were prepped in Laemmli buffer after which 15 - 30 µg of protein was loaded for SDS-PAGE. Antibodies (and their working dilutions) used in the current study are provided in the Supplemental Table 2. Proper loading was initially confirmed by Ponceau S then by probing with anti-β-actin or LMNA antibodies or otherwise indicated. The blot images were captured and analyzed by Odyssey® Fc Imaging System and Image Studio software (LI-COR Biosciences version 5.2), respectively.

### Microscopy and histopathological analysis

H&E and Masson’s Trichrome staining were performed by Translational Research & Pathology Shared Resources (Thomas Jefferson University, PA) using standard methods. Fibrosis quantification of Masson’s Trichrome staining (Fig. 5B) was performed using ImageJ^72^. Briefly, % fibrotic area was derived by dividing area comprised of blue pixels in each micrograph by total number of pixels taken up by the heart tissue in an indicated number of independent images per condition. For immunofluorescence, cells or tissue sections were fixed in 3:1 ratio of ice-cold methanol:acetone, blocked with 10% normal goat serum (ThermoFisher Scientific, cat# 50062Z) and processed using standard methods. Quantitation of immunofluorescence staining of vimentin and PDGFRα (Fig. 5E) were performed similarly to fibrosis quantitation using ImageJ; % positive area was derived by dividing area comprised of the stained pixels (green for vimentin, red for PDGFRα) in each micrograph by total number of pixels. Phalloidin staining was performed using Alexa Fluor® 594 phalloidin (ThermoFisher Scientific, cat# A12381). CFs were fixed on 10% paraformaldehyde followed by formaldehyde-quenching by incubating fixed cells with 0.1M glycine in PBS for 5 min. Phalloidin was diluted 1:100 in PBS and incubated with CFs for 15 min after which they were rinsed 3 times in PBS, counterstained with DAPI, and rehydrated with Prolong^TM^ Diamond Antifade Mountant (ThermoFisher Scientific, cat# P36961). The stained cells were visualized using EVOS™ M7000 Imaging System (ThermoFisher Scientific, cat# AMF7000) with 10X and 20X LED Olympus objective controlled with the EVOS M7000 software. Phalloidin and αSMA quantitation was performed by counting phalloidin and αSMA positive cells divided by the total number of nuclei in the field. Cells displaying phalloidin-stained actin stress fibers traversing the nucleus were considered to be positive. NII was determined using an ImageJ plugin that was previously described^46^. The plugin calculates the percentage of cells that are “irregular”, which are divided into additional subtypes (mitotic, apoptotic, small and irregular, senescent, and large and irregular, and just irregular). We combine and present them as a single “irregular” category without the mitotic subtype (which was typically negligible). Nuclear shape analysis was performed on freshly dissociated CMs that were fixed in 4% paraformaldehyde and stained with DAPI. The CM nuclei images were captured with the Nikon A1R confocal galvano (non-resonant) scanning microscope using a 40X oil immersion objective (Fluor 40X oil, NA1.30, WD 0.20mm) controlled by NIS Elements Imaging Software (version 5.11.02 64bit, Build 1369). Nuclear measurements were performed and calculated on ImageJ using a NDi6d plugin (Nikon ND2 file format reader) by a blinded observer unaware of the genotype.

### Collagen pad contraction assay

Collagen pad contraction assay was performed as previously described^23^. Briefly, CFs at 400,000 cells/ml were combined with 4 mg/ml rat tail type I collagen solution (Advanced Biomatrix, cat# 5153) at a 3:1 ratio to achieve 1 mg/ml collagen I solution containing 3×10^5^ CFs/ml. 500 μl of this solution was aliquoted into each well of a 24-well and allowed to polymerize at 37 °C for 1hr. Following polymerization, the pads were released from the wells and transferred to a 6-well plate with 2 ml of serum-free DMEM, DMEM with 10% FBS, or DMEM with 10% FBS and 10ng/ml TGF-β1 (Prospec, cat# cyt-716). The collagen pads were imaged at 0, 24, and 48 hours and the area of the collagen pads was measured using the circle tool on ImageJ^72^. At the conclusion of the brightfield image capture, the collagen pads were fixed in 4% paraformaldehyde, stained with DAPI, and cell numbers assessed to ensure comparable cell numbers in the collagen pads.

### Scratch migration assay

CFs were grown to ∼90% confluency at which the supplements were gradually removed as described previously. On the following day, after the removal of DMEM with 10% FBS and 10 μM of y-27632, the CFs were rinsed 3X with PBS followed by the addition of fresh DMEM with 0.1% FBS to suppress proliferation. Using a P-200 pipette tip, a scratch line was carefully but firmly made across the confluent monolayer of CFs. The cells were rinsed with PBS to remove the dislocated cells and then incubated at 37°C until imaging at the indicated timepoints. The media was changed daily (DMEM with 0.1% FBS) to remove dead cells. Image capture was performed with Life Technologies EVOS M7000 fluorescence and phase contrast microscope with 10X LED Olympus objective controlled with the EVOS M7000 software.

### M-mode echocardiography

Anesthetized mice (1-2% isoflurane) were placed on a stereotactic heated scanning base (37°C) connected to an electrocardiographic monitor. Left ventricle functional assessment was performed using a Vevo 2100 imaging system (VisualSonics) with a MS550D transducer (22–55 MHz). Parameters were captured for at least three cardiac cycles between the heart rate of 400 to 500 beats per minute. A minimum of three independent M-mode images were used to obtain the cardiac cycle parameters. An echocardiographer, blind to mouse genotype and/or treatment groups, performed the measurements and interpreted the results using AutoLV Analysis Software (VisualSonics).

### Survival analysis

Survival analysis was conducted until time of death or signs of overt distress requiring euthanasia. Mice were assigned to their experimental groups by simple randomization based on sex and littermate information. Specific signs of distress included 1) difficulty with normal ambulation, 2) inability to eat or drink, 3) weight loss of more than 20%, 4) depression, 5) unkempt hair coat, and 6) severe respiratory distress. A staff veterinarian in the Office of Animal Resources at Thomas Jefferson University, blind to experimental groups, determined whether euthanasia was required. Those requiring euthanasia were sacrificed according to the protocol of the Institute of Comparative Medicine consistent with American Veterinary Medical Association guidelines.

### Statistical analysis

Prism 10 (GraphPad) was used to perform statistical analyses. Statistical significance of binary comparisons was determined by unpaired, 2-tailed Student’s *t-*test with *p* ≤ 0.05 considered significant. Statistical significance of three or more comparisons, in which mean values were compared to all other means, was determined by one-way ANOVA with post-hoc Tukey error correction for multiple comparisons. For statistical significance in which three or more means were compared to a defined control, one way ANOVA with Dunnett’s multiple comparison test was used. For statistical significance in which three or more means with 2 or more dependent variables were compared to each other, two-way ANOVA with Tukey correction was used. Statistical significance of median survival from the Kaplan-Meier plot was derived by using a log-rank (Mantel-Cox) test. Survival analysis sample sizes are denoted in the Fig. legends. Values with error bars shown in Fig.s indicate means ± SEM unless stated otherwise. P ≤ 0.05 was considered significant.

## Supporting information

Suppl Figs and Tables combined

## Author Contributions

KS, EP, and JCC conceptualized the research. KS, EP, NB, DM, NW, performed the experiments. GS provided *Lmna*^H222P/H222P^ mice as well as advice and interpretation of experiments. KBM supervised experiments relating to MREs. JCC wrote the original manuscript. All other authors contributed to the review and editing of the final version of the manuscript.

## Acknowledgments

We thank Drs. Raymond Penn, Howard J. Worman, and Qing Chen for insightful discussions. We thank Drs. Yuexing Yuan and Elham Javed for assisting with AutoLV/echo measurements and confocal microscopy analysis, respectively.

## References

1. Wong X, Melendez-Perez AJ, Reddy KL. The Nuclear Lamina. Cold Spring Harbor perspectives in biology. 2022;14. doi: 10.1101/cshperspect.a040113

2. Lin F, Worman HJ. Structural organization of the human gene encoding nuclear lamin A and nuclear lamin C. The Journal of biological chemistry. 1993;268:16321–16326.

3. Dauer WT, Worman HJ. The nuclear envelope as a signaling node in development and disease. Developmental cell. 2009;17:626–638. doi: 10.1016/j.devcel.2009.10.016

4. Fatkin D, MacRae C, Sasaki T, Wolff MR, Porcu M, Frenneaux M, Atherton J, Vidaillet HJ, Jr., Spudich S, De Girolami U, et al. Missense mutations in the rod domain of the lamin A/C gene as causes of dilated cardiomyopathy and conduction-system disease. N Engl J Med. 1999;341:1715–1724. doi: 10.1056/NEJM199912023412302

5. Brayson D, Shanahan CM. Current insights into LMNA cardiomyopathies: Existing models and missing LINCs. Nucleus. 2017;8:17–33. doi: 10.1080/19491034.2016.1260798

6. Taylor MR, Fain PR, Sinagra G, Robinson ML, Robertson AD, Carniel E, Di Lenarda A, Bohlmeyer TJ, Ferguson DA, Brodsky GL, et al. Natural history of dilated cardiomyopathy due to lamin A/C gene mutations. J Am Coll Cardiol. 2003;41:771–780. doi: S0735109702029546 [pii]

7. van Tintelen JP, Tio RA, Kerstjens-Frederikse WS, van Berlo JH, Boven LG, Suurmeijer AJ, White SJ, den Dunnen JT, te Meerman GJ, Vos YJ, et al. Severe myocardial fibrosis caused by a deletion of the 5’ end of the lamin A/C gene. J Am Coll Cardiol. 2007;49:2430-2439. doi: 10.1016/j.jacc.2007.02.063

8. McNally EM, Mestroni L. Dilated Cardiomyopathy: Genetic Determinants and Mechanisms. Circ Res. 2017;121:731–748. doi: 10.1161/CIRCRESAHA.116.309396

9. Watkins H, Ashrafian H, Redwood C. Inherited cardiomyopathies. N Engl J Med. 2011;364:1643-1656. doi: 10.1056/NEJMra0902923

10. Hasselberg NE, Haland TF, Saberniak J, Brekke PH, Berge KE, Leren TP, Edvardsen T, Haugaa KH. Lamin A/C cardiomyopathy: young onset, high penetrance, and frequent need for heart transplantation. Eur Heart J. 2018;39:853–860. doi: 10.1093/eurheartj/ehx596

11. Kumar S, Baldinger SH, Gandjbakhch E, Maury P, Sellal JM, Androulakis AF, Waintraub X, Charron P, Rollin A, Richard P, et al. Long-Term Arrhythmic and Nonarrhythmic Outcomes of Lamin A/C Mutation Carriers. J Am Coll Cardiol. 2016;68:2299–2307. doi: 10.1016/j.jacc.2016.08.058

12. van Rijsingen IA, Nannenberg EA, Arbustini E, Elliott PM, Mogensen J, Hermans-van Ast JF, van der Kooi AJ, van Tintelen JP, van den Berg MP, Grasso M, et al. Gender-specific differences in major cardiac events and mortality in lamin A/C mutation carriers. Eur J Heart Fail. 2013;15:376–384. doi: 10.1093/eurjhf/hfs191

13. Hoorntje ET, Bollen IA, Barge-Schaapveld DQ, van Tienen FH, Te Meerman GJ, Jansweijer JA, van Essen AJ, Volders PG, Constantinescu AA, van den Akker PC, et al. Lamin A/C-Related Cardiac Disease: Late Onset With a Variable and Mild Phenotype in a Large Cohort of Patients With the Lamin A/C p.(Arg331Gln) Founder Mutation. Circ Cardiovasc Genet. 2017;10. doi: 10.1161/CIRCGENETICS.116.001631

14. Kato K, Takahashi N, Fujii Y, Umehara A, Nishiuchi S, Makiyama T, Ohno S, Horie M. LMNA cardiomyopathy detected in Japanese arrhythmogenic right ventricular cardiomyopathy cohort. J Cardiol. 2016;68:346–351. doi: 10.1016/j.jjcc.2015.10.013

15. Rober RA, Weber K, Osborn M. Differential timing of nuclear lamin A/C expression in the various organs of the mouse embryo and the young animal: a developmental study. Development. 1989;105:365–378.

16. Auguste G, Rouhi L, Matkovich SJ, Coarfa C, Robertson MJ, Czernuszewicz G, Gurha P, Marian AJ. BET bromodomain inhibition attenuates cardiac phenotype in myocyte-specific lamin A/C-deficient mice. The Journal of clinical investigation. 2020. doi: 10.1172/JCI135922

17. Chai RJ, Werner H, Li PY, Lee YL, Nyein KT, Solovei I, Luu TDA, Sharma B, Navasankari R, Maric M, et al. Disrupting the LINC complex by AAV mediated gene transduction prevents progression of Lamin induced cardiomyopathy. Nat Commun. 2021;12:4722. doi: 10.1038/s41467-021-24849-4

18. En A, Bogireddi H, Thomas B, Stutzman AV, Ikegami S, LaForest B, Almakki O, Pytel P, Moskowitz IP, Ikegami K. Pervasive nuclear envelope ruptures precede ECM signaling and disease onset without activating cGAS-STING in Lamin-cardiomyopathy mice. Cell Rep. 2024;43:114284. doi: 10.1016/j.celrep.2024.114284

19. Tan CY, Wong JX, Chan PS, Tan H, Liao D, Chen W, Tan LW, Ackers-Johnson M, Wakimoto H, Seidman JG, et al. Yin Yang 1 Suppresses Dilated Cardiomyopathy and Cardiac Fibrosis Through Regulation of Bmp7 and Ctgf. Circ Res. 2019;125:834–846. doi: 10.1161/CIRCRESAHA.119.314794

20. Sikder K, Phillips E, Zhong Z, Wang N, Saunders J, Mothy D, Kossenkov A, Schneider T, Nichtova Z, Csordas G, et al. Perinuclear damage from nuclear envelope deterioration elicits stress responses that contribute to LMNA cardiomyopathy. Sci Adv. 2024;10:eadh0798. doi: 10.1126/sciadv.adh0798

21. Wang J, Chen H, Seth A, McCulloch CA. Mechanical force regulation of myofibroblast differentiation in cardiac fibroblasts. Am J Physiol Heart Circ Physiol. 2003;285:H1871–1881. doi: 10.1152/ajpheart.00387.2003

22. Santiago JJ, Dangerfield AL, Rattan SG, Bathe KL, Cunnington RH, Raizman JE, Bedosky KM, Freed DH, Kardami E, Dixon IM. Cardiac fibroblast to myofibroblast differentiation in vivo and in vitro: expression of focal adhesion components in neonatal and adult rat ventricular myofibroblasts. Dev Dyn. 2010;239:1573–1584. doi: 10.1002/dvdy.22280

23. Shinde AV, Humeres C, Frangogiannis NG. The role of alpha-smooth muscle actin in fibroblast-mediated matrix contraction and remodeling. Biochim Biophys Acta Mol Basis Dis. 2017;1863:298–309. doi: 10.1016/j.bbadis.2016.11.006

24. Kanisicak O, Khalil H, Ivey MJ, Karch J, Maliken BD, Correll RN, Brody MJ, SC JL, Aronow BJ, Tallquist MD, et al. Genetic lineage tracing defines myofibroblast origin and function in the injured heart. Nat Commun. 2016;7:12260. doi: 10.1038/ncomms12260

25. Zheng B, Zhang Z, Black CM, de Crombrugghe B, Denton CP. Ligand-dependent genetic recombination in fibroblasts: a potentially powerful technique for investigating gene function in fibrosis. The American journal of pathology. 2002;160:1609–1617. doi: 10.1016/S0002-9440(10)61108-X

26. Hinz B. The extracellular matrix and transforming growth factor-beta1: Tale of a strained relationship. Matrix Biol. 2015;47:54–65. doi: 10.1016/j.matbio.2015.05.006

27. Small EM, Thatcher JE, Sutherland LB, Kinoshita H, Gerard RD, Richardson JA, Dimaio JM, Sadek H, Kuwahara K, Olson EN. Myocardin-related transcription factor-a controls myofibroblast activation and fibrosis in response to myocardial infarction. Circ Res. 2010;107:294–304. doi: 10.1161/CIRCRESAHA.110.223172

28. Travers JG, Kamal FA, Robbins J, Yutzey KE, Blaxall BC. Cardiac Fibrosis: The Fibroblast Awakens. Circ Res. 2016;118:1021–1040. doi: 10.1161/CIRCRESAHA.115.306565

29. Davis J, Molkentin JD. Myofibroblasts: trust your heart and let fate decide. J Mol Cell Cardiol. 2014;70:9–18. doi: 10.1016/j.yjmcc.2013.10.019

30. Hinz B, Phan SH, Thannickal VJ, Galli A, Bochaton-Piallat ML, Gabbiani G. The myofibroblast: one function, multiple origins. The American journal of pathology. 2007;170:1807–1816. doi: 10.2353/ajpath.2007.070112

31. Driesen RB, Nagaraju CK, Abi-Char J, Coenen T, Lijnen PJ, Fagard RH, Sipido KR, Petrov VV. Reversible and irreversible differentiation of cardiac fibroblasts. Cardiovascular research. 2014;101:411–422. doi: 10.1093/cvr/cvt338

32. Hurley MM, Abreu C, Harrison JR, Lichtler AC, Raisz LG, Kream BE. Basic fibroblast growth factor inhibits type I collagen gene expression in osteoblastic MC3T3-E1 cells. The Journal of biological chemistry. 1993;268:5588–5593.

33. Burger PE, Dowdle EB, Lukey PT, Wilson EL. Basic fibroblast growth factor antagonizes transforming growth factor beta-mediated erythroid differentiation in K562 cells. Blood. 1994;83:1808–1812.

34. Kennedy SH, Qin H, Lin L, Tan EM. Basic fibroblast growth factor regulates type I collagen and collagenase gene expression in human smooth muscle cells. The American journal of pathology. 1995;146:764–771.

35. Ichiki Y, Smith EA, LeRoy EC, Trojanowska M. Basic fibroblast growth factor inhibits basal and transforming growth factor-beta induced collagen alpha 2(I) gene expression in scleroderma and normal fibroblasts. J Rheumatol. 1997;24:90–95.

36. Fang MA, Glackin CA, Sadhu A, McDougall S. Transcriptional regulation of alpha 2(I) collagen gene expression by fibroblast growth factor-2 in MC3T3-E1 osteoblast-like cells. J Cell Biochem. 2001;80:550–559.

37. Papetti M, Shujath J, Riley KN, Herman IM. FGF-2 antagonizes the TGF-beta1-mediated induction of pericyte alpha-smooth muscle actin expression: a role for myf-5 and Smad-mediated signaling pathways. Invest Ophthalmol Vis Sci. 2003;44:4994–5005.

38. Kawai-Kowase K, Sato H, Oyama Y, Kanai H, Sato M, Doi H, Kurabayashi M. Basic fibroblast growth factor antagonizes transforming growth factor-beta1-induced smooth muscle gene expression through extracellular signal-regulated kinase 1/2 signaling pathway activation. Arterioscler Thromb Vasc Biol. 2004;24:1384–1390. doi: 10.1161/01.ATV.0000136548.17816.07

39. Kraemer PM, Ray FA, Brothman AR, Bartholdi MF, Cram LS. Spontaneous immortalization rate of cultured Chinese hamster cells. J Natl Cancer Inst. 1986;76:703–709.

40. Shay JW, Wright WE, Werbin H. Defining the molecular mechanisms of human cell immortalization. Biochim Biophys Acta. 1991;1072:1–7.

41. Arimura T, Helbling-Leclerc A, Massart C, Varnous S, Niel F, Lacene E, Fromes Y, Toussaint M, Mura AM, Keller DI, et al. Mouse model carrying H222P-Lmna mutation develops muscular dystrophy and dilated cardiomyopathy similar to human striated muscle laminopathies. Human molecular genetics. 2005;14:155–169. doi: ddi017 [pii]10.1093/hmg/ddi017

42. Chang W, Antoku S, Ostlund C, Worman HJ, Gundersen GG. Linker of nucleoskeleton and cytoskeleton (LINC) complex-mediated actin-dependent nuclear positioning orients centrosomes in migrating myoblasts. Nucleus. 2015;6:77–88. doi: 10.1080/19491034.2015.1004947

43. Folker ES, Ostlund C, Luxton GW, Worman HJ, Gundersen GG. Lamin A variants that cause striated muscle disease are defective in anchoring transmembrane actin-associated nuclear lines for nuclear movement. Proc Natl Acad Sci U S A. 2011;108:131–136. doi: 10.1073/pnas.1000824108

44. Harada T, Swift J, Irianto J, Shin JW, Spinler KR, Athirasala A, Diegmiller R, Dingal PC, Ivanovska IL, Discher DE. Nuclear lamin stiffness is a barrier to 3D migration, but softness can limit survival. J Cell Biol. 2014;204:669–682. doi: 10.1083/jcb.201308029

45. Khatau SB, Bloom RJ, Bajpai S, Razafsky D, Zang S, Giri A, Wu PH, Marchand J, Celedon A, Hale CM, et al. The distinct roles of the nucleus and nucleus-cytoskeleton connections in three-dimensional cell migration. Sci Rep. 2012;2:488. doi: 10.1038/srep00488

46. Filippi-Chiela EC, Oliveira MM, Jurkovski B, Callegari-Jacques SM, da Silva VD, Lenz G. Nuclear morphometric analysis (NMA): screening of senescence, apoptosis and nuclear irregularities. PloS one. 2012;7:e42522. doi: 10.1371/journal.pone.0042522

47. Bashour AM, Bloom GS. 58K, a microtubule-binding Golgi protein, is a formiminotransferase cyclodeaminase. The Journal of biological chemistry. 1998;273:19612–19617. doi: 10.1074/jbc.273.31.19612

48. Kovacs MT, Vallette M, Wiertsema P, Dingli F, Loew D, Nader GPF, Piel M, Ceccaldi R. DNA damage induces nuclear envelope rupture through ATR-mediated phosphorylation of lamin A/C. Mol Cell. 2023;83:3659–3668 e3610. doi: 10.1016/j.molcel.2023.09.023

49. Pfeifer CR, Xia Y, Zhu K, Liu D, Irianto J, Garcia VMM, Millan LMS, Niese B, Harding S, Deviri D, et al. Constricted migration increases DNA damage and independently represses cell cycle. Molecular biology of the cell. 2018;29:1948–1962. doi: 10.1091/mbc.E18-02-0079

50. Xia Y, Ivanovska IL, Zhu K, Smith L, Irianto J, Pfeifer CR, Alvey CM, Ji J, Liu D, Cho S, et al. Nuclear rupture at sites of high curvature compromises retention of DNA repair factors. J Cell Biol. 2018;217:3796–3808. doi: 10.1083/jcb.201711161

51. Corbin EA, Vite A, Peyster EG, Bhoopalam M, Brandimarto J, Wang X, Bennett AI, Clark AT, Cheng X, Turner KT, et al. Tunable and Reversible Substrate Stiffness Reveals a Dynamic Mechanosensitivity of Cardiomyocytes. ACS Appl Mater Interfaces. 2019;11:20603–20614. doi: 10.1021/acsami.9b02446

52. Kaur H, Takefuji M, Ngai CY, Carvalho J, Bayer J, Wietelmann A, Poetsch A, Hoelper S, Conway SJ, Mollmann H, et al. Targeted Ablation of Periostin-Expressing Activated Fibroblasts Prevents Adverse Cardiac Remodeling in Mice. Circ Res. 2016;118:1906–1917. doi: 10.1161/CIRCRESAHA.116.308643

53. Muchir A, Shan J, Bonne G, Lehnart SE, Worman HJ. Inhibition of extracellular signal-regulated kinase signaling to prevent cardiomyopathy caused by mutation in the gene encoding A-type lamins. Human molecular genetics. 2009;18:241–247. doi: 10.1093/hmg/ddn343

54. Nikolova V, Leimena C, McMahon AC, Tan JC, Chandar S, Jogia D, Kesteven SH, Michalicek J, Otway R, Verheyen F, et al. Defects in nuclear structure and function promote dilated cardiomyopathy in lamin A/C-deficient mice. The Journal of clinical investigation. 2004;113:357–369. doi: 10.1172/JCI19448

55. Antoku S, Wu W, Joseph LC, Morrow JP, Worman HJ, Gundersen GG. ERK1/2 Phosphorylation of FHOD Connects Signaling and Nuclear Positioning Alternations in Cardiac Laminopathy. Developmental cell. 2019;51:602–616 e612. doi: 10.1016/j.devcel.2019.10.023

56. Mounkes LC, Kozlov SV, Rottman JN, Stewart CL. Expression of an LMNA-N195K variant of A-type lamins results in cardiac conduction defects and death in mice. Human molecular genetics. 2005;14:2167–2180. doi: 10.1093/hmg/ddi221

57. Sullivan T, Escalante-Alcalde D, Bhatt H, Anver M, Bhat N, Nagashima K, Stewart CL, Burke B. Loss of A-type lamin expression compromises nuclear envelope integrity leading to muscular dystrophy. J Cell Biol. 1999;147:913–920.

58. Jahn D, Schramm S, Schnolzer M, Heilmann CJ, de Koster CG, Schutz W, Benavente R, Alsheimer M. A truncated lamin A in the Lmna -/- mouse line: implications for the understanding of laminopathies. Nucleus. 2012;3:463–474. doi: 10.4161/nucl.21676

59. Rouhi L, Auguste G, Zhou Q, Lombardi R, Olcum M, Pourebrahim K, Cheedipudi SM, Asghar S, Hong K, Robertson MJ, et al. Deletion of the Lmna gene in fibroblasts causes senescence-associated dilated cardiomyopathy by activating the double-stranded DNA damage response and induction of senescence-associated secretory phenotype. J Cardiovasc Aging. 2022;2. doi: 10.20517/jca.2022.14

60. Artus J, Panthier JJ, Hadjantonakis AK. A role for PDGF signaling in expansion of the extra-embryonic endoderm lineage of the mouse blastocyst. Development. 2010;137:3361–3372. doi: 10.1242/dev.050864

61. Kuwabara JT, Hara A, Heckl JR, Pena B, Bhutada S, DeMaris R, Ivey MJ, DeAngelo LP, Liu X, Park J, et al. Regulation of extracellular matrix composition by fibroblasts during perinatal cardiac maturation. J Mol Cell Cardiol. 2022;169:84–95. doi: 10.1016/j.yjmcc.2022.05.003

62. Kuwabara JT, Hara A, Bhutada S, Gojanovich GS, Chen J, Hokutan K, Shettigar V, Lee AY, DeAngelo LP, Heckl JR, et al. Consequences of PDGFRalpha(+) fibroblast reduction in adult murine hearts. eLife. 2022;11. doi: 10.7554/eLife.69854

63. Koenig AL, Shchukina I, Amrute J, Andhey PS, Zaitsev K, Lai L, Bajpai G, Bredemeyer A, Smith G, Jones C, et al. Single-cell transcriptomics reveals cell-type-specific diversification in human heart failure. Nat Cardiovasc Res. 2022;1:263–280. doi: 10.1038/s44161-022-00028-6

64. Li W, Lou X, Zha Y, Qin Y, Zha J, Hong L, Xie Z, Yang S, Wang C, An J, et al. Single-cell RNA-seq of heart reveals intercellular communication drivers of myocardial fibrosis in diabetic cardiomyopathy. eLife. 2023;12. doi: 10.7554/eLife.80479

65. Farbehi N, Patrick R, Dorison A, Xaymardan M, Janbandhu V, Wystub-Lis K, Ho JW, Nordon RE, Harvey RP. Single-cell expression profiling reveals dynamic flux of cardiac stromal, vascular and immune cells in health and injury. eLife. 2019;8. doi: 10.7554/eLife.43882

66. Ruiz-Villalba A, Romero JP, Hernandez SC, Vilas-Zornoza A, Fortelny N, Castro-Labrador L, San Martin-Uriz P, Lorenzo-Vivas E, Garcia-Olloqui P, Palacio M, et al. Single-Cell RNA Sequencing Analysis Reveals a Crucial Role for CTHRC1 (Collagen Triple Helix Repeat Containing 1) Cardiac Fibroblasts After Myocardial Infarction. Circulation. 2020;142:1831–1847. doi: 10.1161/CIRCULATIONAHA.119.044557

67. McLellan MA, Skelly DA, Dona MSI, Squiers GT, Farrugia GE, Gaynor TL, Cohen CD, Pandey R, Diep H, Vinh A, et al. High-Resolution Transcriptomic Profiling of the Heart During Chronic Stress Reveals Cellular Drivers of Cardiac Fibrosis and Hypertrophy. Circulation. 2020;142:1448–1463. doi: 10.1161/CIRCULATIONAHA.119.045115

68. Kim Y, Zheng Y. Generation and characterization of a conditional deletion allele for Lmna in mice. Biochem Biophys Res Commun. 2013;440:8–13. doi: 10.1016/j.bbrc.2013.08.082

69. Louch WE, Sheehan KA, Wolska BM. Methods in cardiomyocyte isolation, culture, and gene transfer. J Mol Cell Cardiol. 2011;51:288–298. doi: 10.1016/j.yjmcc.2011.06.012

70. Stewart SA, Dykxhoorn DM, Palliser D, Mizuno H, Yu EY, An DS, Sabatini DM, Chen IS, Hahn WC, Sharp PA, et al. Lentivirus-delivered stable gene silencing by RNAi in primary cells. RNA. 2003;9:493–501. doi: 10.1261/rna.2192803

71. Livak KJ, Schmittgen TD. Analysis of relative gene expression data using real-time quantitative PCR and the 2(-Delta Delta C(T)) Method. Methods. 2001;25:402–408. doi: 10.1006/meth.2001.1262S1046-2023(01)91262-9 [pii]

72. Abramoff MD, Magalhaes PJ, Ram SJ. Image Processing with ImageJ. Biophotonics International. 2004;11:36–42.

