## Supplementary material for "Cardiac fibroblasts counterbalance cardiomyocytes in *LMNA* cardiomyopathy pathogenesis": Suppl Figs and Tables combined

### Supplemental Figure S1

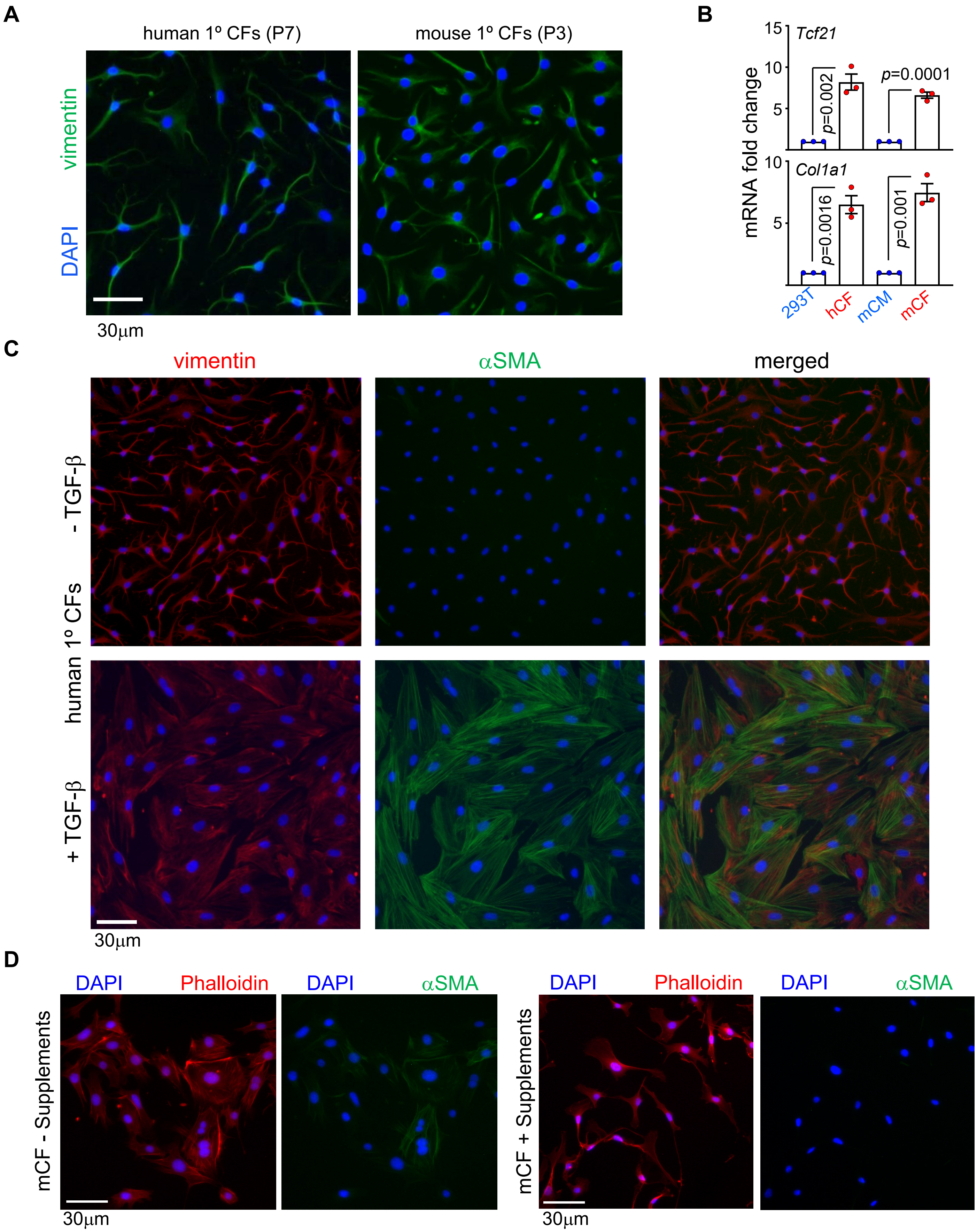


**Supplemental Figure S1. A**) Representative micrographs of hCFs and mCFs stained with anti-vimentin antibodies and DAPI. P7/P3 denote passage number, **B**) qPCR analyses of *Tcf21* and *Col1a1* mRNA in hCFs and mCFs presented as fold change relative to 293T cells and murine CMs, respectively, set to 1. *p* values = unpaired, 2-tailed Student's t-test. n = 3. **C**) Representative micrographs of hCFs with and without 10 ng/ml TGF-β for 96 hr. Untreated cells were maintained with supplements in the culture media. Supplements were gradually removed prior to treatment with TGF-β (see Methods). **D**) Representative micrographs of mCFs cultured in the presence and absence of supplements and stained with anti-SMA antibodies, phalloidin, and DAPI.

Supplemental Figure S2

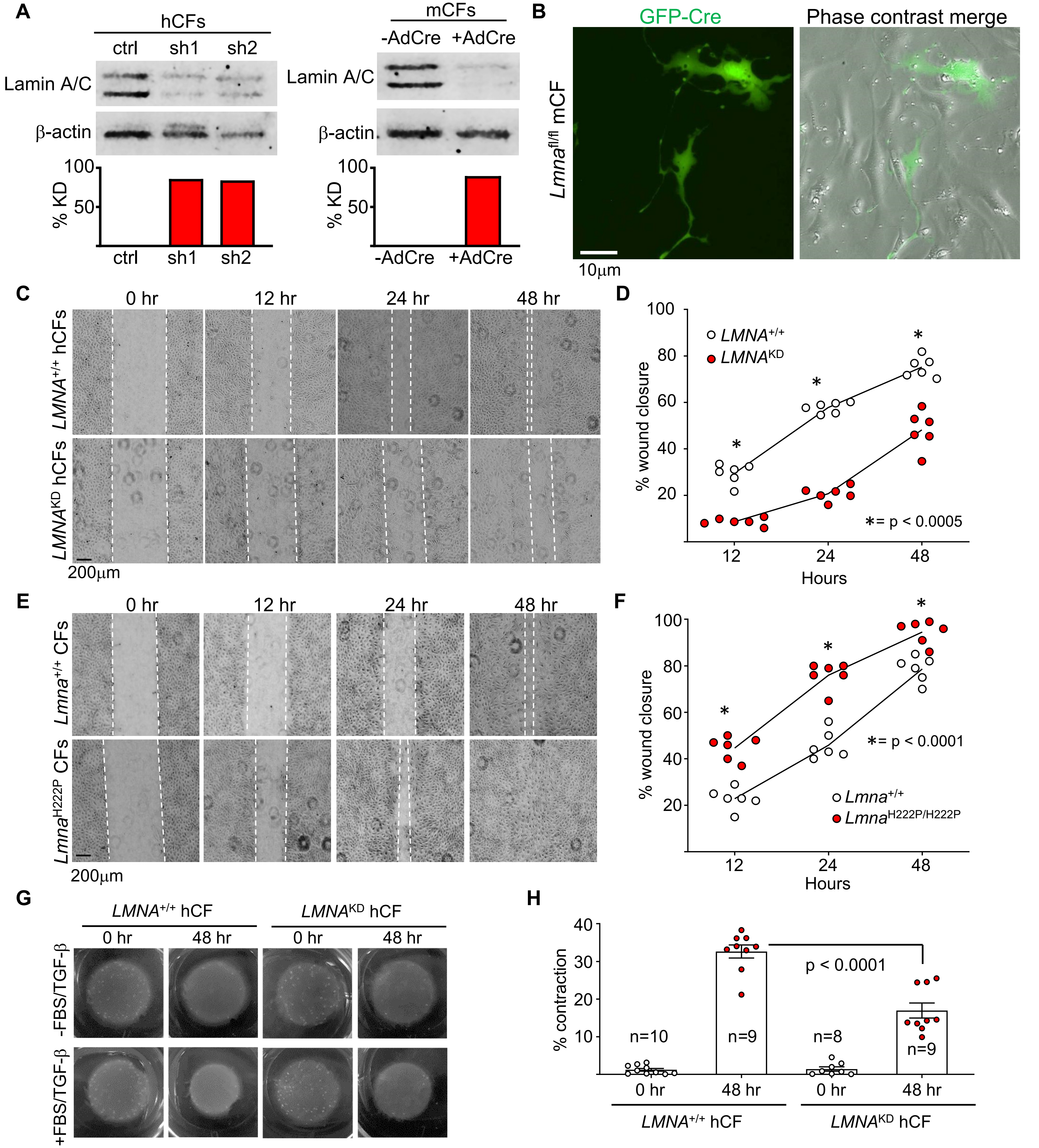


**Supplemental Figure S2. A**) Representative lamin A/C immunoblots on hCFs and mCFs with β-actin as loading control. KD quantitation is shown on the bottom. **B**) Representatiave fluorescence and phase-contrast micrographs of *Lmna*$^{fl/fl}$ mCFs transfected with GFP-Cre. **C-F**) Representative migration micrographs for *LMNA*$^{+/+}$ and *LMNA*$^{KD}$ hCFs (**C**) and their quantitation (**D**) as well as images for *Lmna*$^{+/+}$ and *Lmna*$^{H222P}$ mCFs (**E**) and their quantitation (**F**). n = 3 experiments. *p* values = two-way ANOVA with Bonferroni correction. **G**) Representative collagen pad contraction assay images for *LMNA*$^{+/+}$ and *LMNA*$^{KD}$ hCFs with and without 10% FBS and 10 ng/ml TGF-β. **H**) Results from (**G**) from n = 3 experiments presented as % contraction relative to respective 0 hr. *p* value = one-way ANOVA with Tukey correction. Error bars = SEM.

**Supplemental Figure S3**

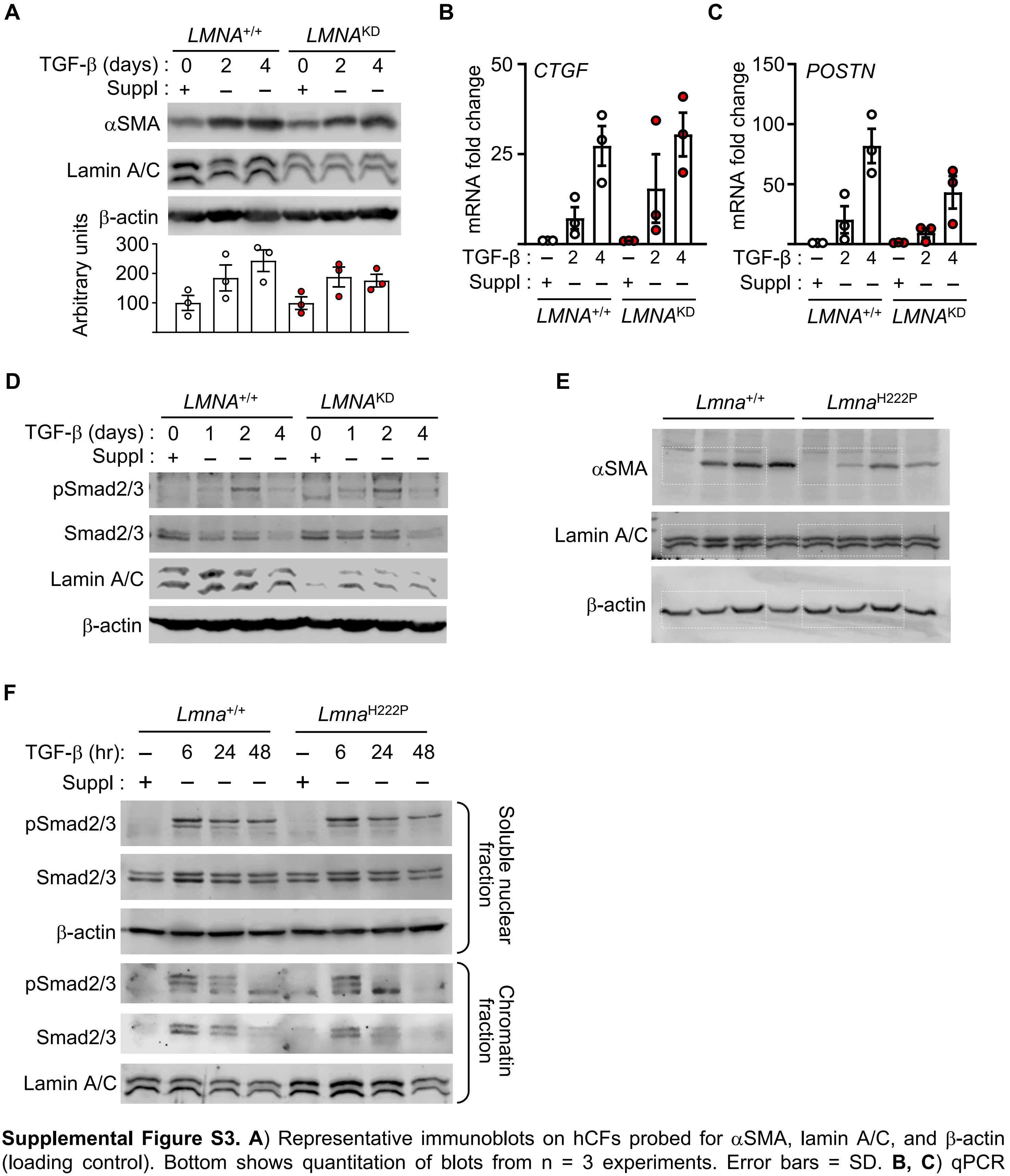


**Supplemental Figure S3. A**) Representative immunoblots on hCFs probed for αSMA, lamin A/C, and β-actin (loading control). Bottom shows quantitation of blots from n = 3 experiments. Error bars = SD. **B, C**) qPCR analyses on samples from (**A**) probed for *CTGF* (**B**) and *POSTN* (**c**) mRNA expression. "-", "2", and "4" denote untreated, 2, and 4 day 10 ng/ml TGF-β treatment, respectively."+", "-", and "Suppl" denote treated or untreated with supplements. Error bars = SEM from 3 experiments. **D**) Representative immunoblots (n = 3) on hCFs probed for Smad2/3 phosphorylation following 10 ng/ml TGF-β stimulation at indicated times after supplement removal. **E**) Uncropped images from Fig 2n showing contiguous blots. **F**) Immunoblots on nuclear fractionation studies on *Lmna*$^{+/+}$ and *Lmna*$^{H222P/H222P}$ mCFs probed for Smad2/3 phosphorylation. β-actin and Lamin A/C were used as loading controls. Representative images from n = 3 experiments are shown.

### Supplemental Figure S4

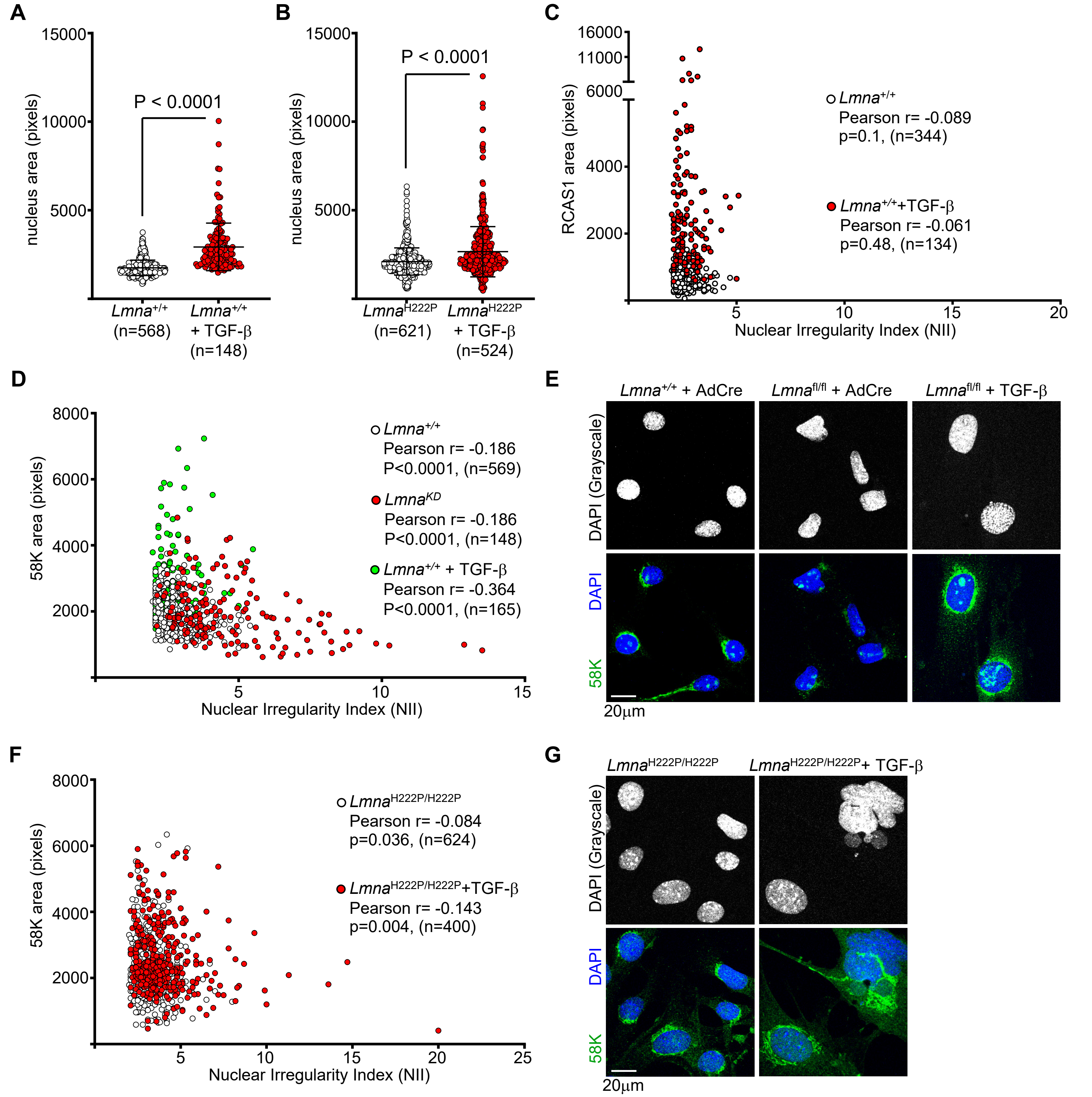


**Supplemental Figure S4. A, B**) Nucleus area of *Lmna*$^{+/+}$ (**A**) and *Lmna*$^{H222P/H222P}$ (**B**) mCFs with and without 10 ng/ml TGF-β. *p* values = unpaired, 2-tailed Student's t-test. Error bars = SD. **C**) NII plotted against perinuclear RCAS1 signal area from *Lmna*$^{+/+}$ mCFs with and without 10 ng/ml TGF-β. **D**) NII plotted against perinuclear 58K signal area from *Lmna*$^{+/+}$, *Lmna*$^{fl/fl}$ + AdCre (*Lmna*$^{KD}$), and *Lmna*$^{+/+}$ + 10 ng/ml TGF-β (48 hr) mCFs. **E**) Representative micrographs of DAPI and 58K stained mCFs from Extended Fig. 4d. **F**) NII plotted against perinuclear 58K signal area from *Lmna*$^{H222P/H222P}$ mCFs with and without 10 ng/ml TGF-β for 48 hr. **G**) Representative micrographs of DAPI and 58K stained mCFs from Supplemental figure S4F.

**Supplemental Figure S5**

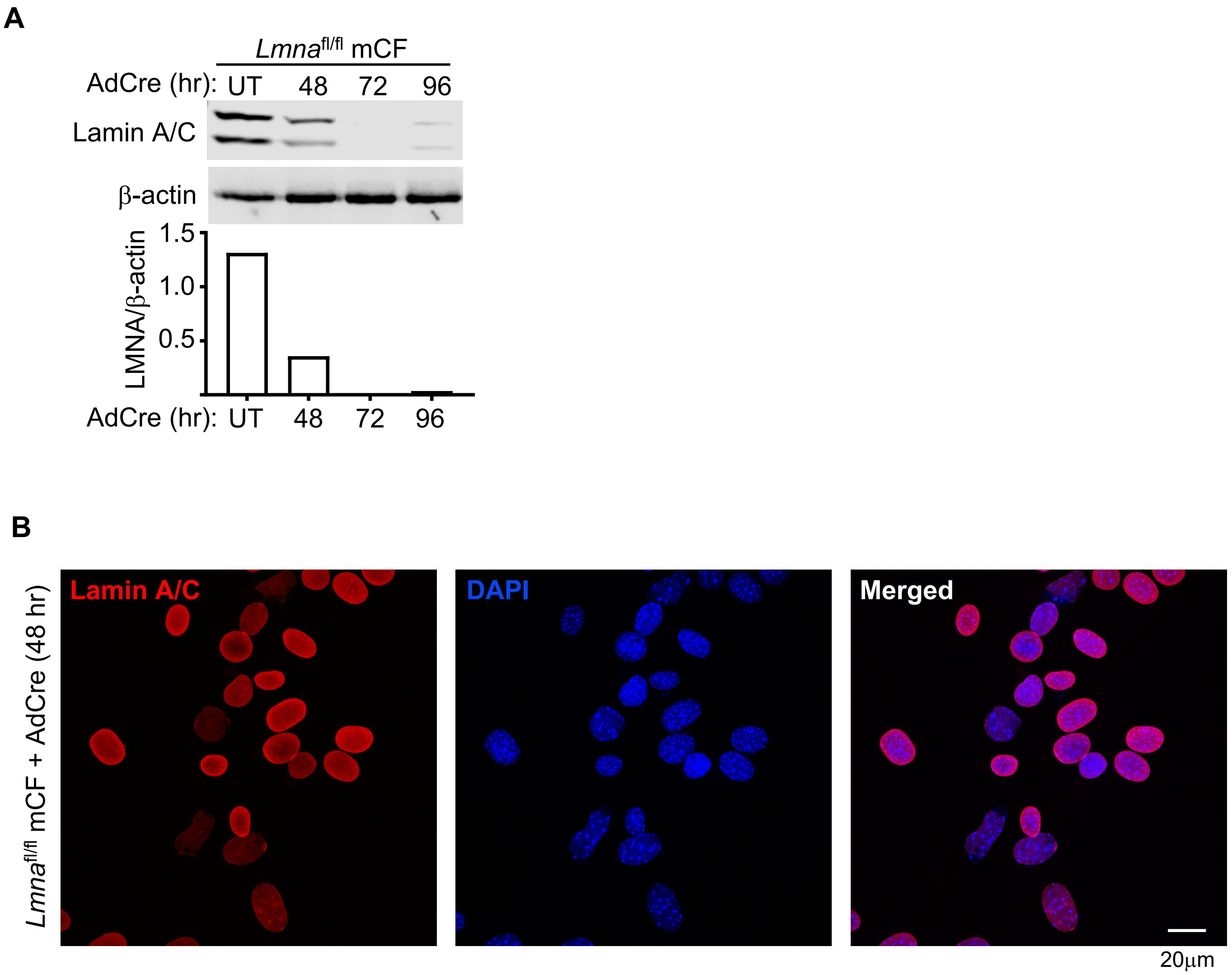


**Supplemental Figure S5. A**) Representative immunoblot of lamin A/C showing KD kinetics after *Lmna*$^{fl/fl}$ mCF infection with AdCre. β-actin was used as loading control. Bottom panel shows quantitation in arbitrary units. **B**) Immunofluorescence micrographs showing Lamin A/C staining with DAPI counterstain on *Lmna*$^{fl/fl}$ mCFs + AdCre for 48 hr.

### Supplemental Figure S6

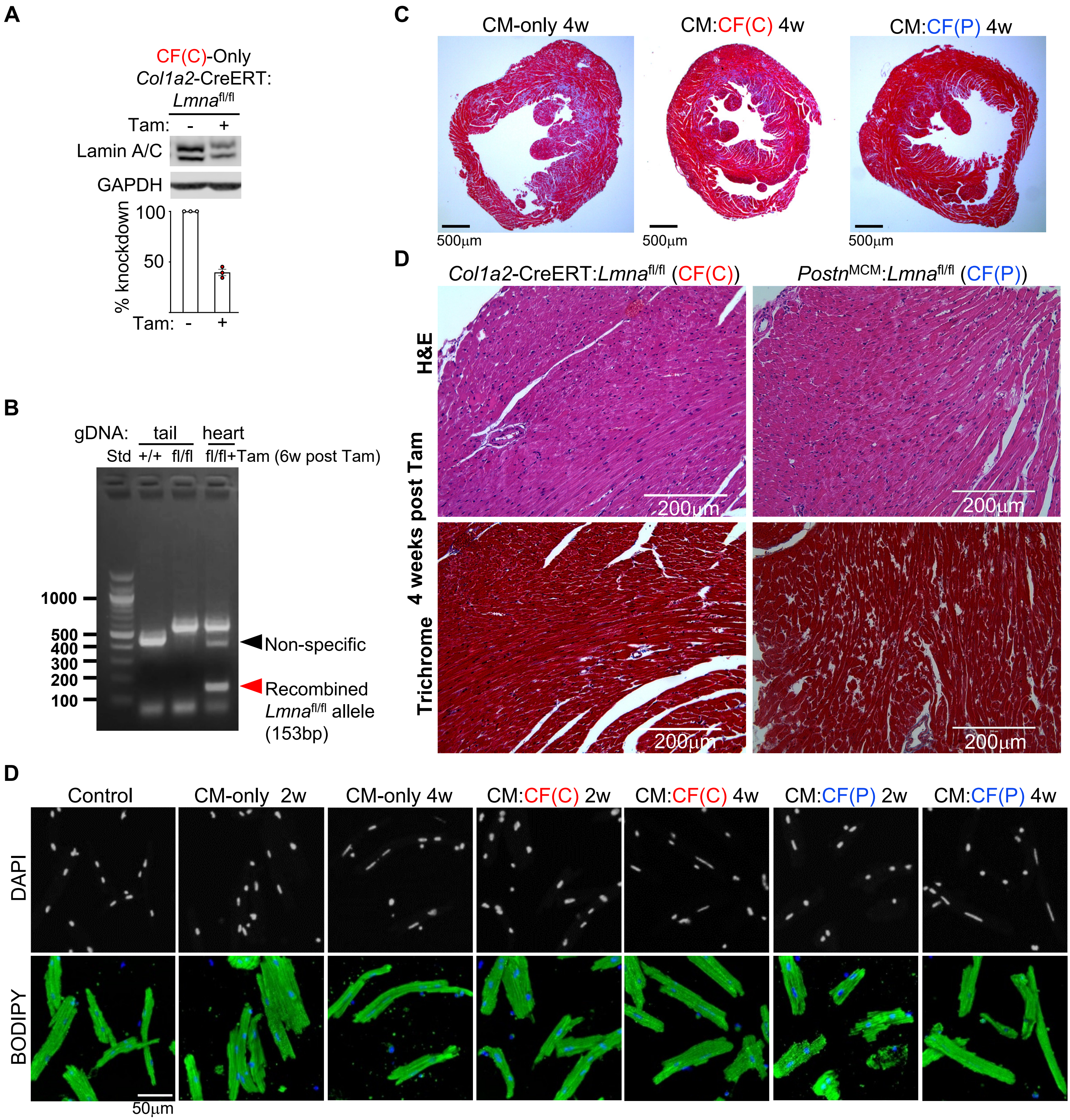


**Supplemental Figure S6. A**) Representative immunoblot of LMNA showing KD in ventricular tissue extract from CF(C)-Only mouse after 4 weeks post Tam. Bottom shows % KD from n = 3 biological replicates calculated relative to vehicle-treated CF(C)-Only hearts set to 100%. Error bars = SD. **B**) PCR analyses showing recombined *Lmna*$^{fl/fl}$ allele from genomic DNA (gDNA) isolated from the tail of wildtype (+/+) and *Lmna*$^{fl/fl}$ (fl/fl) mice as well as from the heart of *Lmna*$^{fl/fl}$ mice at 6 weeks post Tam. "Std" denotes DNA standards. Black arrowhead denotes non-specific band. C) Representative cross section of trichrome stained hearts from CM-only, CM:CF(C), and CM:CF(P) mice at 4 weeks post Tam. **D**) Representative Hematoxylin and Eosin (H&E) and Masson's trichrome staining of hearts from control CF(C) and CF(P) at 4 weeks post Tam are shown. **D**) Representative images of adult CMs isolated from the indicated mouse groups for nuclear shape analyses in Fig. 6D are shown. DAPI in grayscale and adult CMs stained green by BODIPY are shown.

| Primers | Forward (5' - 3') | Reverse (5' - 3') |
|---|---|---|
| *Col1a1* | TTC TCC TGG CAA AGA CGG ACT CAA | AGG AAG CTG AAG TCA TAA CCG CCA |
| *COL1A1* | AGC TGT CTT ATG GCT ATG ATG AGA A | CTT CCC CAT CAT CTC CAT TCT TT |
| *Col1a2* | GGC CCC CTG GTA TGA CTG GCT | CGC CAC GGG GAC CAC GAA TC |
| *Ddr2* | TTC CCT GCC CAG CGA GTC CA | ACC ACT GCA CCC TGA CTC CTC C |
| *Ccn2* (*Ctgf*) | GTG CCA GAA CGC ACA CTG | CCC CGG TTA CAC TCC AAA |
| *CCN2* (*CTGF*) | CCT GGT CCA GAC CAC AGA GT | TGG AGA TTT TGG GAG TAC GG |
| *Postn* | ATG TCA TTG ACC GTG TCC TG | AAG AGC GTG AAG TGA CCA TC |
| *POSTN* | TGC CCT GGT TAT ATG AGA ATG GAA G | GAT GCC CAG AGT GCC ATA AAC A |
| *Tgfb1* | AGC CCG AAG CGG ACT ACT AT | TCC ACA TGT TGC TCC ACA CT |
| *Tgfb2* | CCG GAG GTG ATT TCC ATC TA | GCG GAC GAT TCT GAA GTA GG |
| *Tcf21* | ATG CTG GAC TGT GAC TCC CT | GAG CGG GCT TTT CTT AGT GG |
| *TCF21* | GAG CTC CAA CTG CGA GAA TG | AGG GTG GTC TTG AGT CTG GA |
| *Gapdh* | TGC ACC ACC AAC TGC TTA G | GGA TGC AGG GAT GAT GTT C |
| *GAPDH* | AGG TGG TCT CCT CTG ACT TCA ACA | GAC AAA GTG GTC GTT GAG GGC AAT |
| *Rpl13a* | ATG ACA AGA AAA AGC GGA TG | CTT TTC TGC CTG TTT CCG TA |
| *RPL13A* | AGG TCC TGG TGC TTG ATG GT | TTG ATG CCT TCA CAG CGT AC |
| *Nppa* | TCG TCT TGG CCT TTT GGC T | TCC AGG TGG TCT AGC AGG TTC T |
| *Nppb* | AAG TCC TAG CCA GTC TCC AGA | GAG CTG TCT CTG GGC CAT TTC |
| *Col1a2* | GGC CCC CTG GTA TGA CTG GCT | CGC CAC GGG GAC CAC GAA TC |
| *Col3a1* | GTT CTA GAG GATGGCTGTACTAAACACA | TTG CCT TGC GTG TTT GAT ATT C |
| *Fn1-EDA* | CAG AAA TGA CCA TTG AAG GT | ATG AGT CCT GAC ACA ATC AC |

**Supplemental Table S1. qPCR primer sequences used in the study**

| Antibodies | Company | Catalogue # | Concentration<br>IB = immunoblot<br>IF = immunofluorescence |
|---|---|---|---|
| $\alpha$-smooth muscle actin | Abcam | ab5694 | IB(1:2000), IF(1:300) |
| Vimentin | Cell Signaling Technology | 5741 | IF(1:300) |
| Lamin A/C | Santa Cruz Biotechnology | sc-376248 | IB(1:2000), IF(1:300) |
| phospho-Smad2/3 | Cell Signaling Technology | 8828 | IB(1:500) |
| Smad2/3 | Cell Signaling Technology | 5678 | IB(1:200) |
| RCAS1 | Proteintech | 66170-1-Ig | IF(1:200) |
| 58K-Golgi | Novus Biologicals | NB600-412SS | IF(1:200) |
| PDGFR$\alpha$ | RnD Systems | AF1062 | IF(1:100) |
| $\beta$-actin | Cell Signaling Technology | 3700 | IB(1:4000) |
| GAPDH | Millipore Sigma | MAB374 | IB(1:5000) |
| Donkey anti-goat 594 | Invitrogen | A-11058 | IF(1:400) |
| Goat anti-mouse 594 | Invitrogen | A-21044 | IF(1:400) |
| Goat anti-rabbit 488 | Invitrogen | A-11034 | IF(1:400) |
| Goat anti-rabbit 594 | Invitrogen | R-37117 | IF(1:8) |
| Licor green mouse | LI-COR Biosciences | 926-32210 | IB(1:5000) |
| Licor green rabbit | LI-COR Biosciences | 926-32211 | IB(1:5000) |
| Licor red mouse | LI-COR Biosciences | 926-68070 | IB(1:5000) |
| Licor red rabbit | LI-COR Biosciences | 926-68071 | IB(1:5000) |

**Supplemental Table S2. Antibodies and their dilutions used in the study.**
